# A Biologically Interpretable Single-Cell Pathology Foundation Model for Molecular Decoding of H&E Images

**DOI:** 10.64898/2026.11.14.766416

**Authors:** Weiqing Chen, Qing Li, Hao Cheng, Pengzhi Zhang, Yuzhou Chang, Xiaoying Wang, Tu N Tran, Chengxuan Li, Yuping Yang, Yiwei Xiao, Xiaohan Zhao, Weidong Wu, Anjun Ma, Ana Mora, Mauricio Rojas, Sadhna Aggarwal, Steven H. Lin, Shengyu Li, Qin Ma, Guangyu Wang

## Abstract

The microscopic structure of human tissues encodes molecular information that is not directly accessible by visual inspection. Current pathology foundation models improve histological interpretation but operate at multicellular resolution. It remains unclear whether routine histology contains structured molecular and cellular information recoverable at single-cell resolution, which biological properties are associated with this recoverability, and where its limits lie. Here, we introduce Loki2, a single-cell pathology foundation model trained on UniSeg, a pan-tissue resource integrating 15 million cells with segmentation, lineage annotations and measured or histology-inferred transcriptomic profiles. We examine the above question across cellular representation, molecular recoverability, and patient-level relevance. Loki2 captures cellular identity and continuous morphological variation, with interpretable latent dimensions linked to physical nuclear phenotypes through DAPI validation. Transcriptomic retrieval and *in silico* protein staining test molecular recoverability, with gene-level analyses revealing the strongest recovery for genes associated with morphologically distinct cellular identities and functions. Then, patient staging and biomarker discovery connect cellular morphology representations to clinical relevance. Together, these findings establish Loki2 as a biologically interpretable single-cell pathology foundation model and provide the opportunities of recovering cellular and molecular information from routine H&E histology at single-cell resolution.

## Introduction

The morphological architecture of human tissues encodes a dense and interpretable record of the molecular processes that drive development, homeostasis, and disease^1-3^. Recent advances in artificial intelligence (AI) have begun to unlock these biological signals, led by histopathology foundation models trained at a gigapixel scale^4-7^. Image-only encoders have shown that large and diverse whole-slide datasets can capture broad tissue and disease structure, while vision–language models linking H&E images with diagnostic reports have introduced semantic reasoning across clinical contexts^8-10^. These modalities provide, at best, indirect access to molecular states that govern cellular identity, function, and fate. Visual–omics models have attempted to bridge morphology and genomics^11,12^, but they rely on spatial transcriptomics platforms whose spot sizes (e.g., ∼55 μm) aggregate heterogeneous mixtures of cells. As a result, existing models learn averaged associations instead of the fundamental relationship between a cell’s visual phenotype and its underlying molecular program. This resolution mismatch limits the ability to resolve cell-intrinsic relationships between morphology and molecular features. Fundamentally, it remains unclear how much cellular and molecular information routine histology encodes at single-cell resolution and which biological states can be recovered from cellular morphology.

Here, we introduce Loki2, a single-cell pathology foundation model designed to address this question by learning transferable representations that link cellular morphology with cell identity and molecular state directly from routine H&E images. Rather than assuming that the transcriptome is uniformly predictable from morphology, Loki2 provides a framework for asking which biological information is recoverable from histology at single-cell resolution and how that recoverability varies across molecular states. Loki2 is equipped by the Unified Segmentation (UniSeg) database at its core, comprising approximately 15 million cells across 15 tissue types from 65 studies. In UniSeg, each cell is paired with high-accuracy nucleus segmentation, transcriptomic profiles derived from multi-resolution spatial technologies (including Visium HD, Xenium, and histology-imputed single-cell Visium), and annotations spanning 22 major cell lineages based on a human-in-the-loop AI approach. This unprecedented dataset enables the first large-scale training of a model that jointly embeds segmentation, morphology, cell identity, and molecular state at the native resolution of individual cells.

We use Loki2 to examine this concept across three levels of biological progression from single-cell morphology to molecular recoverability, then to patient-level relevance. First, we evaluated the single-cell morphological representation learned by Loki2 through two complementary analyses. Morphology-based cell typing tests whether the embedding captures discrete cellular identity, while morphological pseudotime examines whether it organizes continuous cell-state variation within a lineage. To interpret the embeddings, we further investigated whether the learned latent structure corresponds to recognizable cellular phenotypes, linking specific dimensions to physical features such as chromatin density and nuclear size, with independent validation using paired H&E–DAPI imaging. Second, we evaluated molecular recoverability at both the transcript and protein levels. Single-cell transcriptomic retrieval tests whether morphology can recover transcriptomic states using external scRNA-seq references. Within this analysis, we quantify gene-specific morphology–molecular relationships to determine which molecular features are captured by the learned representation and are most or least recoverable from histology. Protein-level prediction tests whether molecular recoverability extends beyond transcriptomics to independently measured protein expression. Third, we evaluated patient-level relevance through two complementary analyses. Patient staging tests whether single-cell representations, aggregated through multiple instance learning, capture tissue features associated with disease stage. And biomarker discovery examines whether these representations identify cellular phenotypes and associated molecular features linked to patient outcomes, connecting single-cell morphology–molecular relationships to clinical relevance.

Together, these analyses support a central conclusion: routine H&E contains molecular information that is selectively recoverable at single-cell resolution, and Loki2 preferentially retrieves expression patterns associated with morphologically distinguishable cellular identities and functions. Moreover, Loki2 consistently outperforms 14 state-of-the-art specialist and generalist models. Hence, we now present Loki2 as a transferable and biologically interpretable single-cell pathology foundation model whose latent structure is measurably linked to physical cellular morphology. Beyond supporting multiple analytical tasks, Loki2 provides a biologically grounded framework for determining the cellular and molecular features can be recovered from routine histology. By bridging the long-standing disconnect between tissue morphology and cellular genomics, Loki2 enables new classes of biological discovery and provides a reliable foundation for molecular diagnostics in routine clinical workflows.

## Results

### Loki2: A biologically grounded foundation model for single-cell pathology

To enable the training of a foundation model capable of resolving cellular states from H&E images, we constructed UniSeg, a multimodal resource of unprecedented scale and fidelity. UniSeg comprises approximately 15 million cells paired with expert-validated segmentation masks, cell type annotations, and matched spatial transcriptomic profiles (**Fig. 1a**; **Supplementary Table 1, 2**). The dataset integrates data from 65 studies across 15 tissue types, covering diverse pathological states and staining conditions. To ensure single-cell molecular precision across all platforms, we utilized high-resolution ground truth from Visium HD and Xenium, while applying Thor^13^ to generate histology-inferred single-cell profiles for spot-level Visium data (**Extended Data Fig. 1, 2**). This harmonization yielded a unified training dataset spanning 22 major cell lineages (**Fig. 1b**). Cell masks and phenotypic labels were generated through an automated deep learning pipeline that synthesizes histological and molecular signals, followed by rigorous, systematic quality control and expert validation to ensure high annotation fidelity (**Methods**). Importantly, the strength of UniSeg does not lie simply in total cell count, but in the combination of scale, diversity-aware curation, and high-quality paired morphology–molecular supervision. This is particularly important for pathology foundation model training, where preserving representative cellular heterogeneity together with reliable image and annotation quality is more informative than maximizing raw cell numbers alone.

**Fig. 1:**
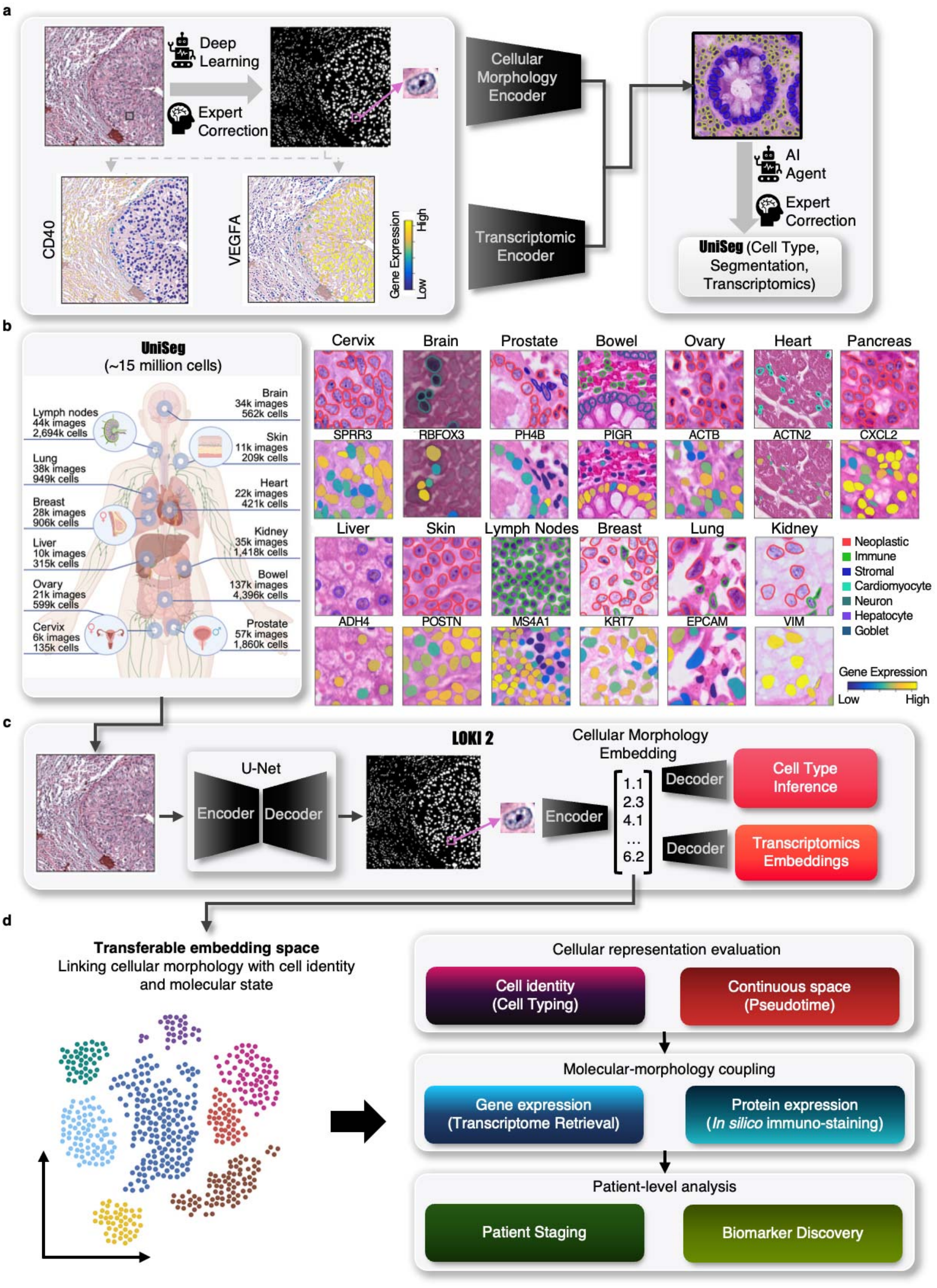
Overview of the study. **a**, The workflow of UniSeg resource preparation. **b**, The left diagram illustrates the size of UniSeg resource in different organs. The right diagram shows the example image patches with cell boundary, type annotation, and paired marker gene expressions from UniSeg. **c**, The workflow of Loki2 model training. **d**, Overview of the Loki2 framework.

Built upon UniSeg, we developed a biologically grounded foundation model, Loki2, to reduce the ambiguity of visual phenotypes using molecular supervision. In contrast to pathology foundation models that rely solely on self-supervision or clinical report text^9,14^, Loki2 adopts a molecular-to-morphology knowledge distillation strategy within a encoder–decoder architecture (**Fig. 1c**). The model comprises four coordinated components. First, a U-Net-based^15^ segmentation module performs instance-level detection, isolating individual nuclei within complex tissue architectures. Second, a vision transformer (ViT)^16^ morphology encoder processes a cropped field of view around each cell, generating high-dimensional embeddings that capture nuclear shape, chromatin texture, and cytoplasmic structure. These morphology embeddings are then used by two parallel decoders during training: a cell type decoder, which maps visual features to transcriptome-defined identities, and a transcriptomic embedding decoder, which reconstructs each cell’s position in a universal gene-expression latent space defined by OmiCLIP^11^ (**Methods**). A contrastive learning objective aligns morphology-derived embeddings with these molecular targets, encouraging the encoder to represent aspects of the cell’s underlying biological state. Through this, Loki2 produces histology-based embeddings enriched with latent molecular semantics, providing a transferable representation that link cellular morphology with molecular state and cell identity from histopathology images.

We use Loki2 to test what routine H&E histology reveals about cellular identity and state, which molecular features it can recover, and how this information relates to patient outcomes (**Fig. 1d**). First, cellular representation evaluation tests whether the learned embedding captures cell identity through cell type annotation and organizes continuous cellular states through morphological pseudotime inference. Second, molecular–morphology coupling examines the molecular information recoverable from cellular morphology. We assess this through single-cell transcriptome retrieval, which maps H&E-segmented cells to reference scRNA-seq atlases, and *in silico* immunostaining, which predicts protein-level biomarker expression. Third, patient-level analysis assesses the clinical relevance of these representations through patient staging and biomarker discovery, using cell-level multiple instance learning to connect cellular features with patient-level outcomes. Together, these analyses evaluate how morphological representations encode cellular identity and state, recover associated molecular information, and inform clinically relevant tissue phenotypes.

### Loki2 resolves individual cells and recovers cellular identity across tissues from H&E image

We first examined whether Loki2 can recover cell identity directly from H&E morphology through cell segmentation and cell typing. To evaluate this performance, we first assembled a held-out test set of UniSeg, spanning all 15 tissue types, capturing diverse histological structures and disease states. We benchmarked Loki2 against two task-specific state-of-the-art models, HoVer-Net^17^ and CellViT^18^, both widely used for nuclear segmentation and classification. Performance was assessed using multi-class panoptic quality (mPQ)^19^ and multi-class F1-score (mF1), metrics that jointly evaluate boundary delineation and cell-type assignment. Across all tissues, Loki2 achieved higher mPQ and mF1-scores than both baselines (**Fig. 2a, b**; **Extended Data Fig. 3a, b**), with statistically significant improvements (*P*-value < 0.001, one-sided Mann– Whitney U test). Notably, whereas HoVer-Net and CellViT exhibited marked performance drops on tissue types underrepresented in public datasets (e.g., liver, small intestine, heart, and brain), Loki2 maintained stable accuracy, suggesting that morphology–transcriptomics pretraining enhances generalization beyond settings covered by standard fully supervised approaches.

To further assess generalization, we performed two out-of-distribution evaluations using slide-level and patient-level holdouts. First, we tested Loki2 on 7 held-out WSIs from seven organ types that were excluded from training (**Extended Data Fig. 4a, b**). These slides showed substantial variation in staining intensity, contrast, section quality, and tissue preservation. Loki2 consistently outperformed CellViT and HoVer-Net across all held-out slides (**Extended Data Fig. 4b**). Second, we performed a patient-level holdout evaluation using 8 patients from 8 independent studies spanning heart, skin, colon, kidney, and lung (**Extended Data Fig. 4c**). Loki2 again outperformed CellViT and HoVer-Net across all held-out patients in cell typing. We further applied Loki2 to a dorsolateral prefrontal cortex (DLPFC) section^20^ that was not included in the training set(**Extended Data Fig. 5a**). We segmented all nuclei in the section and extracted single-cell morphology features for each cell. We then quantified the average cellular morphology similarity across annotated anatomical regions and constructed a morphology-based similarity tree (**Extended Data Fig. 5b**). The resulting tree was consistent with cortical cytoarchitecture, with adjacent cortical layers showed greater similarity than more distant layers, while white matter formed the most distinct branch. Layer 1 was also relatively separated from the deeper cortical layers, consistent with its lower cellular density and neuropil-rich architecture. These results indicate that Loki2 captures reproducible morphology-based organization in unseen samples, supporting its generalization ability beyond the training set. Together, these analyses support Loki2’s generalization beyond the in-distribution benchmark.

We further evaluated Loki2 against generalist biomedical foundation model BiomedParse^21^. Performance was quantified using weighted Dice (joint segmentation and typing) and binary Dice (segmentation only). BiomedParse showed improved performance but was limited by its coarse semantic vocabulary and inability to resolve fine-grained cell lineages. Even when considering segmentation accuracy alone, Loki2 achieved higher binary Dice scores and pixel accuracy than all generalist baseline (**Fig. 2c**; **Extended Data Fig. 3c**). Qualitative inspection (**Fig. 2b; Extended Data Fig. 3b**) further supports these observations: generalist models captured broad regions of interest but frequently merged adjacent nuclei or mis-segmented dense clusters, whereas Loki2 more reliably resolved individual nuclear boundaries. These results indicate that while generalist models have wide applicability, specialized architectures such as Loki2 are required for precise single-cell pathology tasks. We also conducted a benchmark with pathology foundation models, in which UNI^7^, GigaPath^5^, and PLIP^10^ were used as frozen feature extractors. For each model, we extracted image features from cell-level image patches and trained a random forest (RF) classifier on the resulting embeddings to predict cell types. Under this evaluation, Loki2 achieved a weighted F1 score of 0.806 and weighted accuracy of 0.807, whereas the competing backbones achieved weighted F1 scores between 0.574 and 0.608 and weighted accuracies between 0.594 and 0.625 (**Extended Data Fig. 6**).

**Fig. 2:**
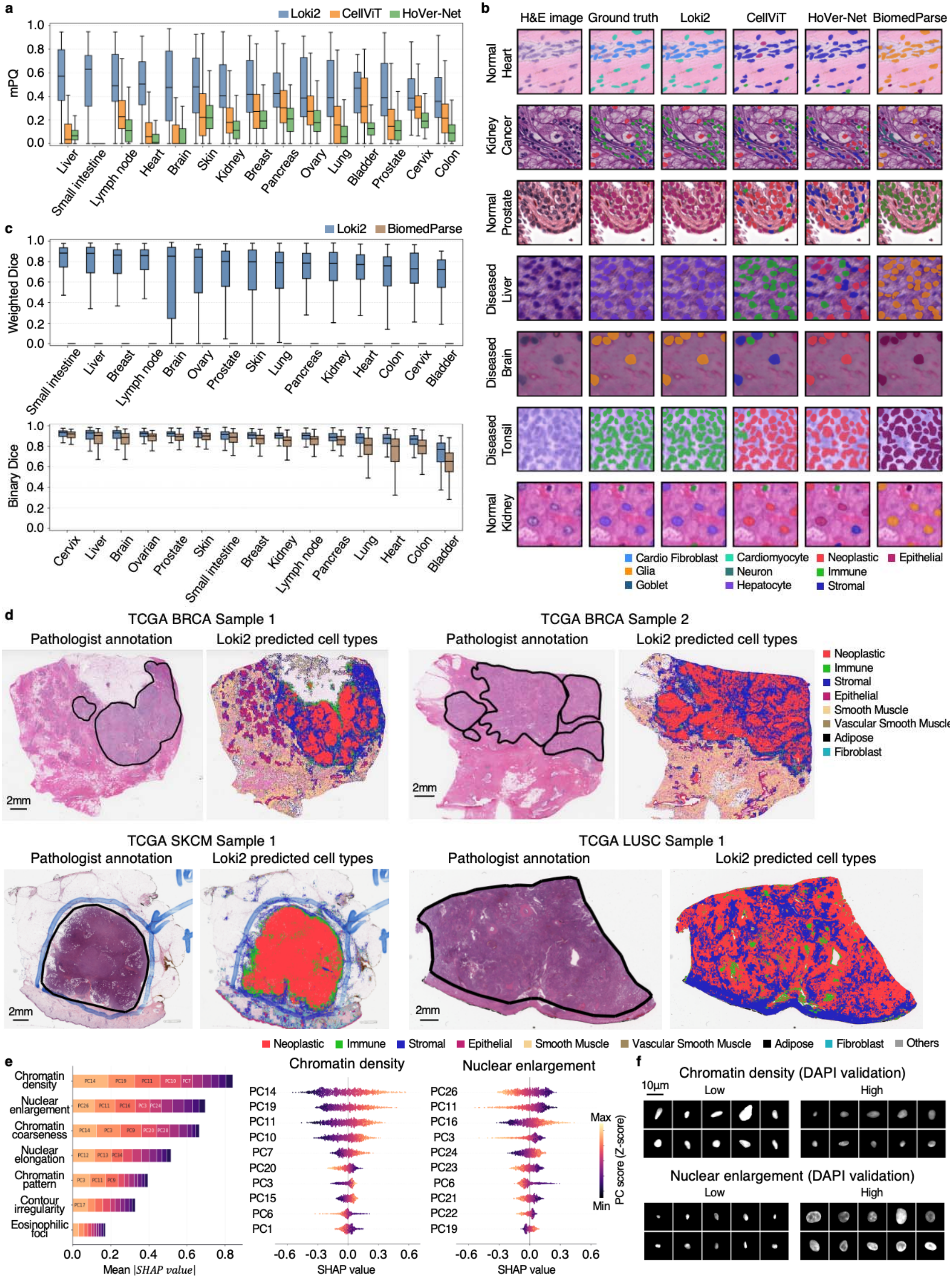
Benchmarking Loki2 for single-cell segmentation, typing, and generalization. **a**, Box plots comparing mPQ scores for Loki2, CellViT, and HoVer-Net across 9,573 test instances from 15 tissue types (sample sizes: colon 2,552; lung 1,300; heart 1,253; prostate 1,000; kidney 753; breast 500; brain 500; skin 400; lymph node 389; liver 250; ovarian 200; small intestine 150; pancreatic 150; bladder 126; cervix 50). In the box plots, the middle line represents the median, the box boundaries indicate the interquartile range, and the whiskers extend to data points within 1.5 times the interquartile range. **b**, Representative examples comparing segmentation and cell-type annotation results from Loki2, CellViT (remapped labels), HoVer-Net (remapped labels), and BiomedParse with ground truth. **c**, Box plots of weighted Dice and binary Dice scores for Loki2 and BiomedParse on the same test instances. Box plot elements follow the conventions described in (c). **d**, Four TCGA examples illustrating Loki2 cell segmentation and annotation alongside pathologist-annotated tumor regions. **e**, SHAP analysis of the seven morphology features predicted from H&E images of 20 held-out samples. Left, mean |SHAP| value for each of seven morphology features, with stacked segments showing the contributing PCs. Middle and right, SHAP beeswarm plots for chromatin density and nuclear enlargement. Each point is one cell; color shows the z-scored PC value. **f**, Representative DAPI nuclei with low and high predicted chromatin density and nuclear enlargement.

To assess robustness in independent clinical settings, we applied Loki2 to H&E-stained whole-slide images (WSIs) from The Cancer Genome Atlas (TCGA)^22^, including breast invasive carcinoma (BRCA), cutaneous melanoma (SKCM), lung squamous cell carcinoma (LUSC), thyroid carcinoma (THCA), and kidney renal clear cell carcinoma (KIRC). We compared Loki2’s single-cell predictions with pathologist-delineated tumor regions. Across samples, we observed close correspondence between model-derived neoplastic cell distributions and expert annotations (**Fig. 2d**; **Extended Data Fig. 3d**). Neoplastic cells predicted by Loki2 were localized within annotated tumor regions, while stromal and immune cells populated the surrounding microenvironment, consistent with expected histological organization. In the melanoma WSI, Loki2 distinguished the central tumor mass from the surrounding capsule and adipose tissue without slide-level fine-tuning, demonstrating its ability to generalize external datasets and diverse tissue presentations.

Finally, to demonstrate population-level utility, we quantified the cellular landscape of HER2-positive breast cancer across 73 TCGA BRCA WSIs. By aggregating Loki2’s single-cell predictions across millions of cells, we constructed population-level cellular composition profiles (**Extended Data Fig. 3e**). These profiles revealed a characteristic cellular architecture with a predominant neoplastic compartment (mean proportion 65.9%), accompanied by variable immune infiltration (mean proportion 20.5%) and stromal content (mean proportion 9.9%).

Importantly, these morphology-derived estimates were consistent with cellular compositions reported in recent molecular benchmarking studies^23^, indicating that Loki2 can recover key features of the tumor microenvironment directly from H&E images. These results support that single-cell morphology encodes recoverable information about cellular identity, enabling characterization of tumor composition and microenvironmental organization directly from routine H&E histology.

### Loki2 embeddings capture interpretable and measurable cellular morphology

To determine whether Loki2 embeddings encode interpretable cellular morphology, we analyzed seven histological phenotypes routinely assessed in computational pathology^24-30^ using a UniSeg discovery cohort comprising 80 training and 20 held-out samples, with no patient overlap between the two sets (**Supplementary Table 3**). SHAP analysis^31^ linked these phenotypes to distinct combinations of embedding principal components (PCs). Chromatin density, nuclear enlargement and chromatin coarseness showed the largest summed mean absolute SHAP values across their ten leading PCs, whereas contour irregularity and eosinophilic foci showed the smallest. Chromatin density was associated primarily with PC14 and PC19, and nuclear enlargement with PC26 and PC11 (**Fig. 2e**).

We next tested the transferability of these associations in an independent cohort of 18 esophageal cancer specimens that differed in tissue type, acquisition site and staining and scanning protocols (**Extended Data Fig. 7a, 8**). The phenotype-associated PCs and predictive relationships identified in the discovery cohort were applied without refitting or feature reselection. SHAP analysis in this cohort recovered a similar phenotype ranking and largely overlapping leading PCs (**Extended Data Fig. 7b**). Within each slide, cells in the upper and lower quartiles of the summed phenotype-specific SHAP scores differed in independently measured H&E morphology in the expected direction. Phenotypic separations ranged from 0.3 to 1.7 within-slide standard deviations in the esophageal cohort, compared with 0.6 to 1.9 in the held-out UniSeg samples (**Extended Data Fig. 7c**), supporting the transferability of the latent-morphology relationships.

To establish whether these relationships correspond to physical nuclear properties measurable beyond H&E, we acquired paired DAPI images of the same nuclei in the 18 esophageal specimens. Cells were stratified within each slide using phenotype-specific SHAP scores derived exclusively from their H&E embeddings, and the corresponding phenotypes were measured independently in DAPI images. Groups defined by chromatin-density-associated PCs differed in DAPI-measured chromatin density, whereas those defined by nuclear-enlargement-associated PCs differed in nuclear size (**Fig. 2f**). The phenotype distributions shifted broadly between the upper and lower quartiles, with separations of 0.4–1.2 within-slide standard deviations across the measurements tested; all but one comparison were significant (*P*-value < 0.01, two-sided Mann–Whitney test; **Extended Data Fig. 7c**). These within-specimen associations provide orthogonal evidence that Loki2 latent structure reflects measurable nuclear morphology.

Finally, we assessed whether seven conventional morphological descriptors could account for the information encoded by Loki2. Cell-type classification using Loki2 embeddings achieved an accuracy and weighted F1 score of 0.80, compared with 0.41 for both metrics using the conventional descriptors (**Extended Data Fig. 7d**). Thus, Loki2 captures information relevant to cellular identity beyond this predefined feature set. Together, these analyses link the learned representation to interpretable, physically measurable cellular phenotypes while demonstrating additional morphological information relevant to cell-type discrimination.

### Morphological pseudotime reveals continuous cell-state transitions

We next asked whether Loki2’s morphological representations capture continuous cell-state variation within a lineage, extending beyond discrete cellular identity. Biological processes such as differentiation, immune activation, and malignant transformation progress through continua of intermediate cell states rather than discrete categories. In single-cell genomics, trajectory inference methods have enabled reconstruction of these dynamic processes by modeling gene-expression changes along a continuous pseudotime^32^. We hypothesized that nuclear morphology undergoes analogous gradual transitions during cell state changes, and that these dynamics are captured within the latent space learned by Loki2. To explore this possibility, we adapted the Markov chain diffusion approach originally developed for transcriptomic analyses (e.g., Palantir^33^). Instead of operating on gene count matrices, our implementation builds a *k*-nearest neighbor (*k*-NN) graph directly from Loki2’s single-cell morphology embeddings. Using biologically motivated root states (such as stem-like or normal epithelial cells), we simulate a Markov random walk over this morphological manifold. The resulting diffusion probabilities define a morphological pseudotime, providing an ordering of cells along developmental or pathological trajectories derived solely from progressive changes in their visual features. To further evaluate the embedding space severing as cellular morphology manifold, we generated low-dimensional space across tissues and observed distinct manifold geometries in the brain and skin (**Methods**; **Extended Data Fig. 9a**). These results suggest that Loki2 embeddings are not arbitrarily distributed in latent space, but instead exhibit structured, tissue-dependent organization consistent with gradual phenotypic variation. The geometry was not uniform across tissues, with linearity scores of 0.84 in skin and 0.55 in brain (**Methods**). We also observed a highly similar low-dimensional manifold structure in the diffusion map (**Extended Data Fig. 9b**) of two independent CRC patients, indicating that the organization is reproducible across patients rather than idiosyncratic to one specimen. We also performed a cross-tissue single-cell morphology–transcriptome similarity analysis across eight tissues (**Extended Data Fig. 10a**), directly quantifying how strongly nuclear morphology reflects transcriptomic state across different biological contexts (**Extended Data Fig. 10b**). This analysis showed substantial tissue-dependent variation, with stronger morphology–molecular coupling in some tissues and weaker coupling in others.

Tumor evolution arises through the gradual accumulation of somatic mutations, which is typically reconstructed using genomic sequencing to infer clonal phylogenies. We hypothesized that this underlying genetic divergence would be reflected in a corresponding morphological continuum within the Loki2 latent space. To investigate this, we analyzed a multi-regional CRC dataset for which the clonal phylogenetic structure had been established using whole-exome sequencing (WES)^34^. The genomic analysis identified an ancestral clone (G1) and a descendant subclone (G2) that further diversified into spatially distinct regions. For each region, we first applied Loki2 to segment the WSI and infer cell-level identities, enabling us to isolate only tumor cells for downstream trajectory analysis. This restriction ensures that the inferred morphological continuum reflects variation within the malignant compartment rather than broader tissue-level heterogeneity. Using these tumor-cell embeddings, we constructed a morphology-based diffusion map across the three sampled regions (G1-clone1, G2-clone2, and G2-clone3), designating the G1 clone as the trajectory root based on its ancestral genomic status (**Extended Data Fig. 11a**). The resulting morphological pseudotime ordering revealed a progression from G1 toward the two G2 subclones (**Fig. 3a**), with G2-clone2 occupying intermediate pseudotime values and G2-clone3 enriched at later positions. This ordering was observed consistently in both the high-dimensional UMAP and projection embeddings (**Fig. 3b**) and the spatial coordinates of the tissue sections. Notably, the pseudotime, inferred from Loki2’s morphological embeddings, was highly consistent with the mutational phylogeny derived from WES, effectively recapitulating the evolutionary relationships among the tumor regions. These observations suggest that subtle morphological features captured by Loki2 reflect aspects of the underlying clonal structure. While not a substitute for genomic sequencing, the alignment between morphological pseudotime and WES-defined clonal relationships indicates that certain phenotypic signatures of genomic divergence may be detectable directly from H&E images, providing a complementary perspective on intratumoral evolutionary dynamics.

**Fig. 3:**
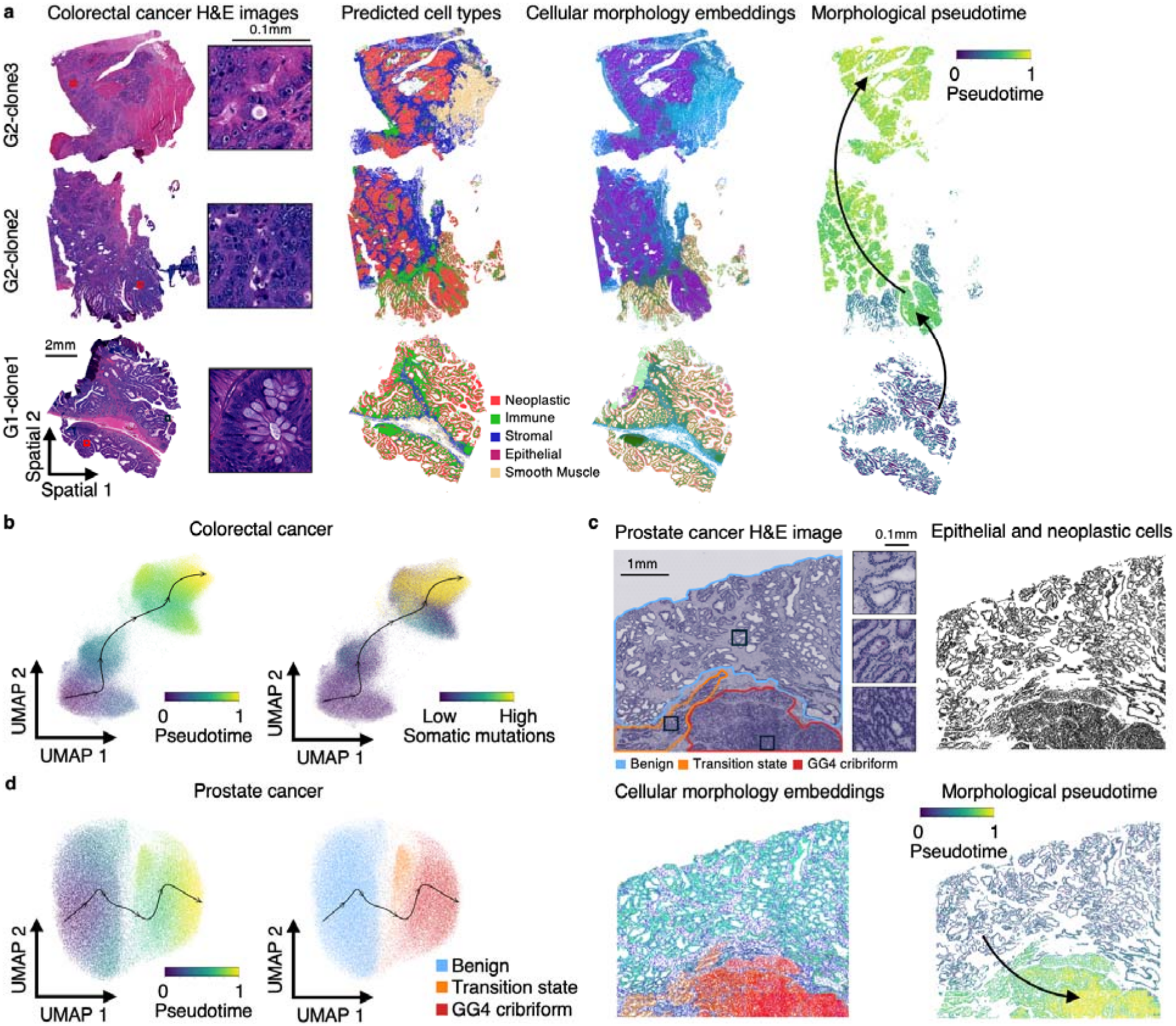
Morphology-based pseudotime inference. **a**, H&E images, Loki2 predicted cell types, cellular morphology embeddings, and inferred morphological pseudotime for three CRC samples. For cellular morphology embeddings, the samples were visualized by mapping their three UMAP components that computed from cellular morphology embeddings to the red, green, and blue channels, respectively. **b**, UMAP visualization of cellular morphology embeddings from the three CRC samples, colored by inferred morphological pseudotime and somatic mutations from the original study, respectively. **c**, H&E image, Loki2 identified epithelial and neoplastic cells, cellular morphology embeddings, and inferred morphological pseudotime for the prostate cancer sample. For cellular morphology embeddings, the samples were visualized by mapping their three UMAP components, which were computed from cellular morphology embeddings to the red, green, and blue channels, respectively. **d**, UMAP visualization of cellular morphology embeddings from the prostate cancer sample, colored by inferred morphological pseudotime and region types from the original study, respectively.

We next investigated whether morphological pseudotime could capture the continuous spectrum of tumor dedifferentiation, using prostate adenocarcinoma as a paradigm. Unlike the clonal branching observed in colorectal cancer, prostate cancer progression is characterized by a gradual loss of glandular organization, from well-formed benign glands to fused or cribriform structures. To examine this process, we analyzed a WSI from a spatially resolved transcriptomics study exhibiting heterogeneous grading patterns^35^ (**Fig. 3c**). Using Loki2, we segmented the tissue and isolated epithelial and neoplastic cells, designating benign glands as the trajectory root (**Extended Data Fig. 11b**). The resulting morphological pseudotime revealed a continuous axis that broadly tracked the histological annotations from the original study. Cells within well-differentiated benign glands occupied early pseudotime values, whereas those within high-grade cribriform regions appeared later along the trajectory. Notably, cells located at the interface between benign and malignant regions were assigned intermediate pseudotime states, capturing transitional morphologies that are often difficult to represent with discrete grading. A similar continuum was visible in the high-dimensional latent space, where cells formed a smooth manifold bridging benign, transition, and malignant compartments, annotated by the original study (**Fig. 3d**). The correspondence between morphological pseudotime and canonical grading patterns suggests that Loki2 captures subtle, gradual remodeling associated with prostate tumor progression, providing a quantitative perspective on dedifferentiation that complements traditional categorical grading.

To further test whether the inferred morphological pseudotime reflects biological insights rather than incidental variation in cell shape or appearance, we coupled pseudotime to transcriptomic programs in the prostate cancer case study. We first clustered genes by their correlation with the inferred pseudotime into three groups: Group 1 (absolute value < 0.2), Group 2 (absolute value between 0.2–0.4), and Group 3 (absolute value > 0.4) (**Extended Data Fig. 12a**). We then performed pathway enrichment analysis within each group to test whether progressively pseudotime-associated genes converged on coherent biological processes. This analysis revealed a clear gradient of biological specificity. Genes in the highly correlated Group 3 showed the strongest and broadest enrichment, mapping to pathways that are highly relevant to prostate cancer progression and cellular state transitions, including DNA repair, androgen response, p53 signaling, and MYC target programs. These genes were also enriched for pathways linked to active cell-state remodeling, such as mTORC1 signaling, unfolded protein response, glycolysis, oxidative phosphorylation, reactive oxygen species pathway activity, fatty acid metabolism, and xenobiotic metabolism (**Extended Data Fig. 12b**). Collectively, these signals indicate coordinated changes spanning proliferation, stress adaptation, oncogenic signaling, and metabolic rewiring. By contrast, the moderately correlated Group 2 showed weaker and more restricted enrichment, whereas the low-correlation Group 1 exhibited limited and less coherent pathway associations overall. This graded enrichment pattern suggests that genes most tightly aligned with morphological pseudotime are also those involved in major tumor-relevant regulatory and metabolic programs, while genes weakly related to pseudotime are less associated with tumor related pathways. Together, these analyses indicate that Loki2 embeddings capture continuous morphological variation associated with clonal relationships, histological differentiation and coordinated molecular programs, extending cellular representation beyond discrete cell identity.

### Loki2 recovers spatial transcriptomic patterns from single-cell morphology

To test whether cellular morphology can support transcriptomic recovery at single-cell resolution, we used Loki2 to retrieve molecular profiles from external scRNA-seq references. This analysis asks whether the morphological representation of an H&E-segmented cell can identify transcriptionally similar cells and recover spatial expression patterns consistent with independently measured molecular data (**Fig. 4a**). To achieve this semantic alignment, we trained a contrastive learning projector using paired high-resolution data from Visium HD, which provides spatially matched morphological and transcriptomic signals. Specifically, the model learns to map the visual embeddings of a segmented nucleus to the corresponding foundation model based transcriptomic embeddings. By optimizing the joint embedding space so that matching image–gene pairs are pulled closer while non-matching pairs are pushed apart, Loki2 establishes a shared latent manifold. Once trained, this model enables zero-shot retrieval: given a cell in an H&E image, the model projects it into this shared space to retrieve the most transcriptionally similar cells from an external scRNA-seq dataset, effectively reconstructing the spatial molecular landscape of the tissue *in silico*.

**Fig. 4:**
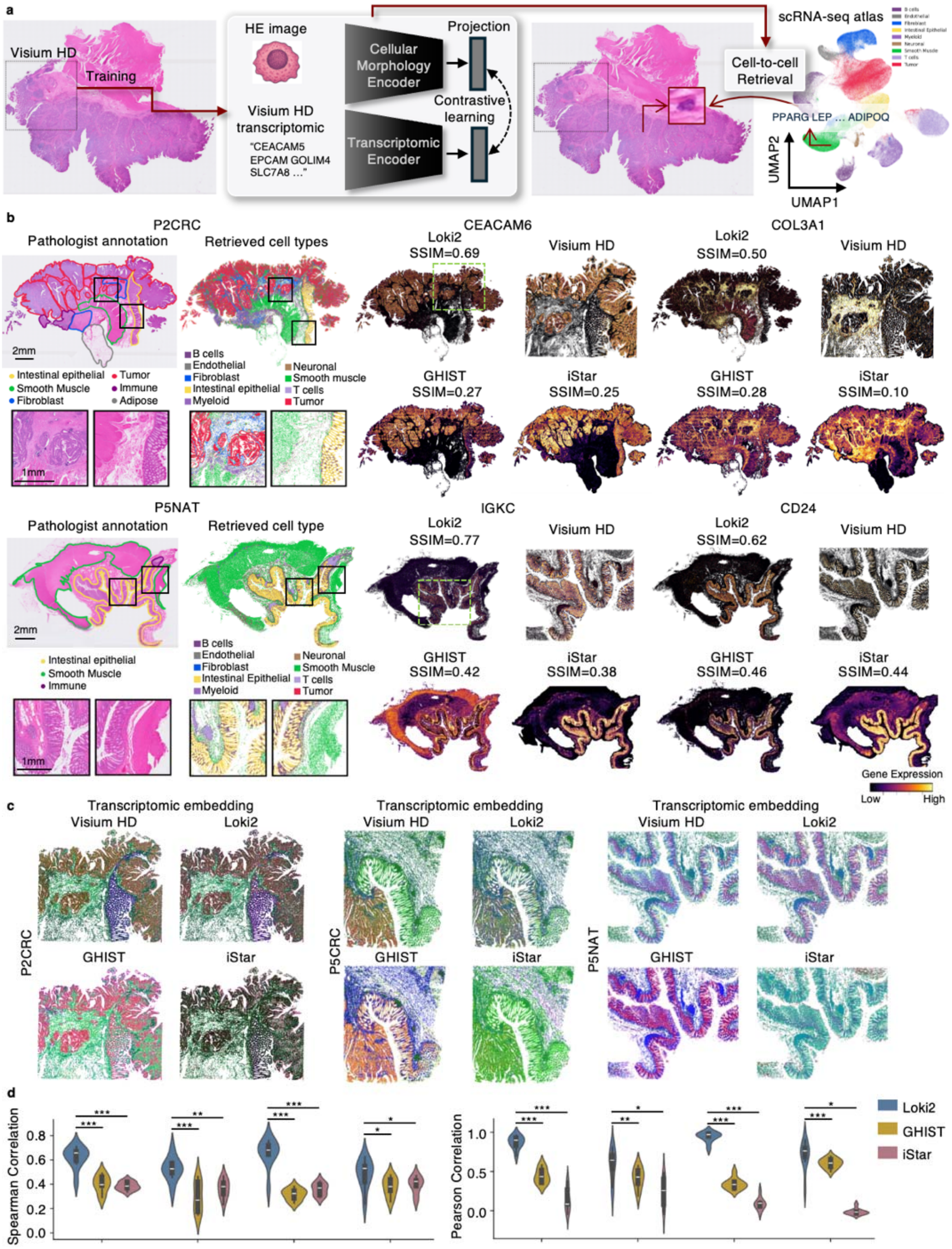
Morphology-to-transcriptome retrieval at single-cell resolution. **a**, Schematic overview of single-cell morphology-to-transcriptome retrieval using Loki2. **b**, Left panels show cell-type annotations from the pathologist and from Loki2, including corresponding zoom-in regions. Right panels show example gene-expression maps from Visium HD (green dashed box region) alongside predictions from Loki2, GHIST, and iStar, with SSIM values computed relative to the Visium HD ground truth. **c**, Visualization of samples by mapping the top three PCA components that fit on Visium HD gene expression and then applied to gene expression predictions from Loki2, GHIST, and iStar, to the red, green, and blue channels, respectively. **d**, Gene expression prediction performance comparison using Spearman correlation and Pearson correlation of highly expressed gene ranks predicted by Loki2, GHIST, and iStar, respectively. Statistical significance was assessed using a one-sided Mann–Whitney U test; \**P*-value □ < 5×10^-2^, \*\**P*-value □< □1×10^-2^, \*\*\**P*-value □< □1×10^-3^, and \*\*\*\**P*-value □ <1×10^-4^. In the box plots, the middle line represents the median, the box boundaries indicate the interquartile range, and the whiskers extend to data points within 1.5 times the interquartile range.

To validate the performance of the retrieval task, we trained the contrastive alignment module using a colorectal cancer (CRC) specimen profiled with Visium HD (**Fig. 4a**), utilizing the platform’s single-cell-scale resolution to establish precise morphology–transcriptomics pairs (**Extended Data Fig. 13a**). We first evaluated the retrieval fidelity by projecting both the H&E-derived morphological embeddings and the ground-truth transcriptomic embeddings into the shared latent space. Visualized via uniform manifold approximation (UMAP) (**Extended Data Fig. 13b, c**), the two modalities exhibited a good overlap (average similarity 0.713), confirming that Loki2 successfully learned a unified manifold where cells with similar visual features map to similar molecular profiles. This alignment was further corroborated spatially: when we projected the morphology-only embeddings back onto the tissue coordinates, they reconstructed a spatial map that was virtually indistinguishable from the ground-truth molecular map (**Extended Data Fig. 13d**). Furthermore, when applying the retrieval module to reconstruct the spatial landscape (**Extended Data Fig. 13e**), Loki2 accurately recovered the complex molecular heterogeneity of the tumor microenvironment, distinguishing tumor nests, stromal bands, and immune niches solely from the H&E input.

To evaluate the generalization capacity of the retrieval module, we applied the trained model to an independent test cohort comprising four colorectal samples from patients excluded from training. To assess robustness to biological heterogeneity, the cohort included both diseased tissue (two colorectal carcinomas) and healthy controls (two normal colon samples), which have H&E WSIs and paired Visium HD as a validation for the Loki2 retrieval task. For each sample, we used the WSI to retrieve nearest-neighbor cells from a colorectal scRNA-seq reference atlas, thereby performing zero-shot spatial transcriptomic reconstruction. We verified the cell type annotation of scRNA-seq reference atlas that canonical marker genes were specifically enriched in their corresponding annotated cell populations (**Extended Data Fig. 14a**). The reconstructed cell-type distributions showed close correspondence with pathologist annotations across both disease states (**Fig. 4b**; **Extended Data Fig. 15**). In carcinoma samples, Loki2 retrieved neoplastic lineages within tumor nests and immune populations within the surrounding stroma, whereas in normal tissue, it recovered the expected epithelial organization of crypt structures.

We further assessed the accuracy of continuous molecular predictions by comparing Loki2’s reconstructed gene expression maps with matched Visium HD data. Performance was benchmarked against GHIST^23^ and iStar^36^, two state-of-the-art (SOTA) spatial gene expression prediction methods. Visually, Loki2 produced structurally coherent expression patterns (e.g., sharp CEACAM6 gradients in epithelial regions and COL3A1 enrichment in stromal bands) while comparison methods frequently yielded over-smoothed or spatially noisy reconstructions. Quantitatively, Loki2 achieved a higher structural similarity index measure (SSIM) for highly expressed genes across all four samples at gene level (*P*-value < 0.001, one-sided Mann– Whitney U test; **Fig. 4b; Extended Data Fig. 15, 16**), indicating improved fidelity in synthesizing spatial transcriptomic landscapes directly from standard histology.

Unlike gene expression prediction methods such as GHIST and iStar, Loki2 achieves higher biological fidelity because each retrieved cell corresponds to an experimentally measured scRNA-seq profile rather than a computationally generated expression vector. This design preserves native gene co-expression structures. To visualize the global preservation of transcriptomic topology, we performed Principal Component Analysis (PCA) on the high-dimensional gene-expression matrix and mapped the first three principal components to RGB color channels^37^ (**Fig. 4c; Extended Data Fig. 17**), enabling a holistic comparison of molecular phenotypes across the tissue. Loki2’s reconstructed landscape closely matched the ground-truth Visium HD data, reproducing the dominant color gradients that delineate major biological compartments. In contrast, GHIST and iStar showed noticeable spectral distortions and reduced structural coherence, often assigning inconsistent molecular signatures to anatomically distinct regions and failing to capture the sharp transitions between tumor nests and adjacent stroma. To evaluate gene-level fidelity, we focused on the top 300 highly expressed genes in each sample and computed both Spearman and Pearson correlation coefficients between the reconstructed and ground-truth expression profiles at cell-type level (**Fig. 4d**). This approach specifically assesses the model’s ability to preserve the relative ordering of gene expression within a tissue, independent of absolute expression magnitude. Across all four patients, Loki2 showed consistently higher rank-based correlations than GHIST and iStar (*P*-value < 0.001, one-sided Mann–Whitney U test). In the P5NAT normal-colon sample, Loki2 achieved a median Pearson correlation near 1.0, indicating close agreement in the reconstructed gene-ordering hierarchy, whereas iStar exhibited correlations near zero in several cases. These results suggest that regression-based methods may be more sensitive to shifts in gene-expression structure, while the retrieval-based approach of Loki2 offers more stable preservation of gene ranking across tissues.

To assess the platform dependency, we performed an experiment using Xenium data from the same colorectal training specimen. All other components of the task were kept unchanged (**Extended Data Fig. 18a**). In carcinoma samples, Loki2 retrieved neoplastic lineages within tumor nests together with immune and stromal populations in the surrounding microenvironment, whereas in normal tissue it recovered the expected epithelial organization of crypt structures (**Extended Data Fig. 18b, 19**). Consistent with the original Visium HD-trained model, the Xenium-trained model generated spatially coherent molecular patterns that aligned closely with tissue architecture and pathologist annotation, such as stromal COL3A1 enrichment and tumor epithelial-associated CEACAM6 expression in representative samples (**Extended Data Fig. 19**). Quantitatively, across all four test samples, the Xenium-trained model achieved performance broadly comparable to the Visium HD-trained model and remained consistently superior to GHIST and iStar in Spearman correlation, Pearson correlation at cell-type level, and SSIM at gene level (**Extended Data Fig. 18c**).

In addition, to assess sensitivity to the choice of reference atlas, we performed the analysis by adding an independent colorectal cancer scRNA-seq atlas from a separate study^38^, which differs in both cohort composition and annotation granularity (**Extended Data Fig. 14b**). Consistent with using the original reference scRNA-seq, the retrieved profiles from independent scRNA-seq atlas also generated spatially coherent molecular patterns that closely aligned with tissue architecture and pathologist annotations (**Extended Data Fig. 20a, 21**). We further constructed variants with different levels of mixing between the original and external atlases to evaluate the stability of downstream results under changing reference conditions (**Extended Data Fig. 20b**). Across these settings, we observed consistent performance in cell type assignment and downstream evaluations, indicating that our conclusions are not dependent on a specific annotation tool or atlas covering major cell types. Moreover, we tested the failure mode of Loki2. Transcriptomic retrieval depended on accurate cell-type matching, with additional, specimen-dependent gains from morphological matching within cell types (**Supplementary Note 1**).

Retrieval is also constrained by reference-atlas compatibility: missing or poorly represented cell types and disease states can lead to mismatched molecular assignments. These findings identify cellular crowding, weak within-cell-type morphology–molecular coupling and incomplete reference coverage as important conditions that can limit prediction accuracy.

Together, these results show that Loki2 can recover morphology-associated transcriptomic patterns in independent colorectal specimens, with consistent performance across two molecular supervision platforms and alternative scRNA-seq references. This supports the recoverability of molecular features coupled to cellular morphology and tissue organization, while retaining dependence on the molecular states represented in the reference atlas.

### Morphology–molecular coupling defines a hierarchy of gene-expression recoverability

Having established that molecular recoverability varies across genes and biological programs, we next examined how this selective coupling translates into spatially resolved transcriptomic retrieval in individual tissues. To define which molecular features are recoverable from single-cell morphology, we examined gene-specific morphology–molecular coupling in 15 Visium HD sections spanning six tissue types, with paired cell-level histological and transcriptomic measurements. For each gene, we calculated the mutual information (MI)^39^ between measured expression and the corresponding Loki2 morphological representation (PCs), then ranked genes into five groups, G1–G5, from highest to lowest MI (**Extended Data Fig. 22a**). High-MI genes showed distinct associations with specific morphological principal components, revealing gene-specific structure within the embedding (**Extended Data Fig. 23**). This hierarchy provided a basis for examining molecular recoverability, biological function and transcriptomic retrieval performance.

We first assessed whether morphology-associated molecular information was predictive through simple linear probes trained to predict individual gene expression from Loki2 embeddings. Linear decoding performance followed the MI hierarchy: well-decoded genes were most frequent in G1 and progressively less frequent towards G5, whereas poorly decoded genes predominated in the lower-MI groups (**Extended Data Fig. 22b**). The concordance between statistical association and linear decoding indicates that genes strongly coupled to the morphological representation also tend to be more readily predicted from it.

Functional analysis revealed biological differences across the recoverability hierarchy (**Extended Data Fig. 22c**; **Methods**). G1 was enriched for specialized secretion, secretory machinery and epithelial differentiation, consistent with established links between these programs and cellular features such as secretory apparatus development and epithelial architecture. G2 combined protein production and quality control with diverse lineage-associated functions. G3 and G4 encompassed immune-associated programs: G3 included immune-cell identity, interferon responses and antigen presentation, whereas G4 emphasized lymphocyte signaling and innate immune responses alongside stromal and vascular functions. Their lower recoverability may reflect functional variation among cells with similar morphological appearances. G5 contained a heterogeneous mixture of cellular maintenance, regulatory and lineage-associated functions, indicating that weak morphology–molecular coupling was not confined to a single biological process. Consistent with these functional differences, mean cell-type specificity decreased from 0.60 in G1 to 0.28 in G5, whereas the mean number of cell types expressing each gene remained broadly similar across groups (2.70–3.03; **Methods**). Highly recoverable genes were therefore more strongly enriched in particular cellular identities, rather than simply expressed in fewer cell types. Gene-level rankings, MI scores and cell-type specificity measures are provided in **Supplementary Table 4**.

We next examined whether the MI hierarchy was reflected in transcriptomic retrieval. Among genes with high retrieval performance (SSIM > 0.3), 48.7% belonged to G1, 36.7% to G2, 13.6% to G3 and 0.9% to G4; none belonged to G5 (**Extended Data Fig. 22d**). Thus, 85.4% of highly retrievable genes fell within the two highest-MI groups, linking successful retrieval to stronger morphology-associated molecular information. Together, these results define a hierarchy of molecular recoverability in the evaluated tissues: Loki2 preferentially recovers expression patterns associated with morphologically distinguishable cellular identities and functions, while molecular features weakly coupled to its morphological representation show more limited recoverability.

### Morphology-associated molecular recoverability extends to protein expression

We next examined whether the nuclear morphology encoded by Loki2 is sufficiently rich to recover protein-level expression directly from standard H&E images. To test this, we developed an *in silico* staining task (**Fig. 5a**) that maps Loki2’s single-cell morphological embeddings to marker-specific protein intensities. We registered the adjacent H&E and multiplexed immunofluorescence (mIF) images, expanded morphological masks, and extracted the marker-level protein intensities. Using datasets containing about 200,000 cells from 12 paired H&E and mIF images, we derived Loki2 embeddings and trained a LightGBM^40^ predictor to reconstruct spatial protein patterns solely from morphology-derived features.

**Fig. 5:**
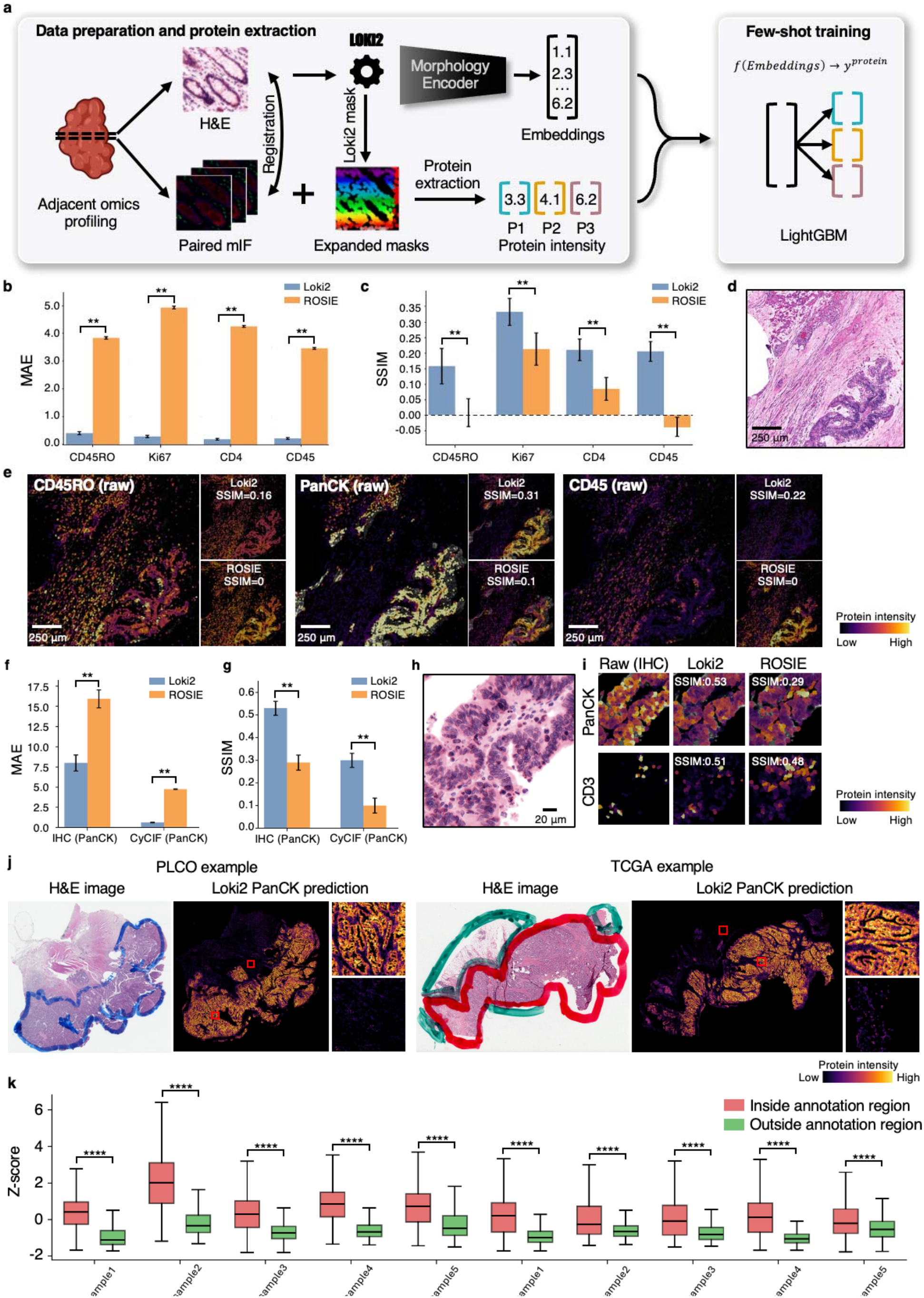
*In silico* immunostaining from standard histology. **a**, The workflow of Loki2 *in silico* immunostaining from standard histology. **b**, Benchmarking results showing MAE across four CyCIF markers, including CD45RO, Ki67, CD4, and CD45. Statistical significance was assessed using a one-sided Mann–Whitney U test; \**P*-value <□ 5×10^-2^, \*\**P*-value□ < □ 1×10^-2^, \*\*\**P*-value□ <□ 1×10^-3^, and \*\*\*\**P*-value□ <1×10^-4^. **c**, Benchmark results showing SSIM values for the same four CyCIF markers. **d**, An example CRC H&E image patch. **e**, Visualization of raw H&E, virtual stain images generated by Loki2 and ROSIE, respectively, across CD45RO, PanCK, and CD45. **f**, Benchmarking results showing MAE for PanCK in both IHC and CyCIF. Statistical significance was assessed using a one-sided Mann–Whitney U test; \**P*-value□ <□ 5×10^-2^, \*\**P*-value □ < □1×10^-2^, \*\*\**P*-value □ < □ 1×10^-3^, and \*\*\*\**P*-value □ <1×10^-4^. **g**, Benchmarking results showing SSIM values for PanCK in IHC and CyCIF. **h**, An example colorectal cancer H&E image patch. **i**, Visualization of the raw H&E image and virtual stain images generated by Loki2 and ROSIE, respectively. **j**, Example whole-slide H&E images and corresponding Loki2-predicted PanCK protein intensity with zoom-in regions from PLCO (left) and TCGA (right) cohorts. **k**, Quantitative comparison of Z-score of Loki2 predicted PanCK intensity inside and outside pathologist annotated tumor regions across multiple samples from PLCO and TCGA cohorts. In the box plots, the middle line represents the median, the box boundaries indicate the interquartile range, and the whiskers extend to data points within 1.5 times the interquartile range.

In a CRC cohort profiled by CyCIF^41^, predictors trained on Loki2 embeddings consistently outperformed the ROSIE model^42^ as well as other baseline prediction models (**Extended Data Fig. 24a**). Loki2 achieved significantly lower mean absolute error (MAE) and higher SSIM across a panel of immune and proliferation markers, including CD45RO, Ki67, CD4, and CD45 (**Fig. 5b, c**; *P*-value < 0.01, one-sided Mann–Whitney U test). Notably, these gains reflect not only improved intensity prediction but also enhanced preservation of spatial architecture. Qualitatively, Loki2 recapitulated complex protein landscapes with high fidelity (**Fig. 5d, e**; **Extended Data Fig. 24b**). In contrast, the baseline model commonly generated blurred or nonspecific patterns in densely cellular regions. Moreover, Loki2 precisely resolved fine-scale boundaries, delineating immune infiltrates and proliferative niches within the heterogeneous tumor microenvironment. To evaluate robustness across staining modalities, we further assessed performance on colon cancer tissues stained by immunohistochemistry (IHC)^43^. Loki2 again demonstrated superior recovery of Pan-Cytokeratin (PanCK) and CD3 distributions relative to all baselines (**Fig. 5f, g**; **Extended Data Fig. 24c**), yielding significant improvements in both MAE and SSIM (*P*-value < 0.01, one-sided Mann–Whitney U test). The accurate reconstruction of epithelial and T-cell compartments across both CyCIF and IHC datasets (**Fig. 5h, i**) indicates that Loki2’s morphological representations generalize across imaging technologies and effectively extract protein-level expression from routine histology.

We further validated the *in silico* immunostaining prediction using external clinical cohorts with pathologist-annotated tumor regions. In this analysis, we focused on PanCK, a well-established epithelial marker widely used in clinical immunohistochemistry. We directly applied the freeze pretrained LightGBM model that previously trained to predict the PanCK protein marker, to the Loki2-derived cell embeddings without any additional fine-tuning, from H&E images of CRC patients across two independent cohorts PLCO and TCGA. The predicted PanCK intensity maps show strong spatial concordance with epithelial tumor regions, with clear enrichment in morphologically defined tumor areas and low signal in surrounding stromal or non-epithelial regions (**Fig. 5j**; **Extended Data Fig. 24d**). To quantitatively validate these predictions, we compared the Z-scored PanCK intensity inside versus outside pathologist-annotated tumor regions across multiple samples from both cohorts. As shown in **Fig. 5k**, the predicted PanCK intensity is consistently and significantly higher within tumor regions than in non-tumor regions. The separation between these two distributions is robust across samples and cohorts, indicating that the model captures biologically meaningful protein expression patterns that generalize beyond the training data. Importantly, this validation leverages independent clinical annotations and external datasets, rather than relying on the training data or internal supervision signals.

Failure mode analysis showed that marker-intensity prediction remained robust to the synthetic segmentation perturbations tested, but performance was more variable in densely packed regions with overlapping nuclei (**Supplementary Note1**). Together, these results extend morphology-associated molecular recoverability to selected protein markers, showing that Loki2 embeddings support prediction of independently measured protein patterns across CyCIF and IHC datasets.

### Cell-level multiple instance learning enables whole-slide clinical prediction

We next asked whether the cellular information encoded by Loki2 is associated with patient-level disease stage. To connect single-cell morphology with whole-slide assessments, we implemented cell-level multiple instance learning (MIL), which aggregates nuclear embeddings across a whole-slide image into a slide-level prediction. This formulation retains individual cells as the units of analysis and assigns attention weights that indicate their relative contribution to the aggregated representation.

To assess whether single-cell morphology provides a sufficient signal for patient-level prognostication, we applied the MIL-based cancer patient staging in TCGA KIRC, bladder urothelial carcinoma (BLCA), and colon adenocarcinoma (COAD) by distinguishing between early and late tumor stages. We compared the Loki2 with PLIP, UNI, and GigaPath’s embedding-based RF classifier. We evaluated the performance by two matrices (AUC and average precision (AP)). Loki2 achieved the top 1 score in every comparison (**Fig. 6a**; **Extended Data Fig. 25**), demonstrating its effectiveness for cancer staging based on cell-level morphology. Specifically, Loki2 achieved AUC 0.91, 0.78, and 0.68 and AP 0.91, 0.75, and 0.66 for KIRC, BLCA, and COAD.

Since the MIL assigns an attention weight to each cell, it provides cell-resolved maps that highlight the elements most influential for stage classification (**Fig. 6b; Extended Data Fig. 26-28**). In three stage tasks, high-attention regions predominantly overlapped with tumor epithelial areas characterized by increased nuclear atypia, cellular crowding, and disrupted architecture, whereas low-attention regions were more frequently located within stromal and immune-rich compartments. These patterns indicate that Loki2 prioritizes epithelial morphology for subtype prediction while assigning lower attention to surrounding non-tumor regions that contribute less discriminatory information in this setting. Beyond recapitulating expected histological distinctions, the resulting attention maps outline the cellular subpopulations enriched within each class, offering a conservative and interpretable view of how different tissue compartments contribute to subtype discrimination. High-attention regions largely coincided with neoplastic epithelial areas (**Fig. 6b; Extended Data Fig. 26-28**). Low-attention regions, in contrast, were more often confined to tissue areas with limited discriminatory value. Together, these analyses connect Loki2’s single-cell morphological representations to patient-level disease stage, supporting the clinical relevance of cellular phenotypes captured from routine H&E histology.

**Fig. 6:**
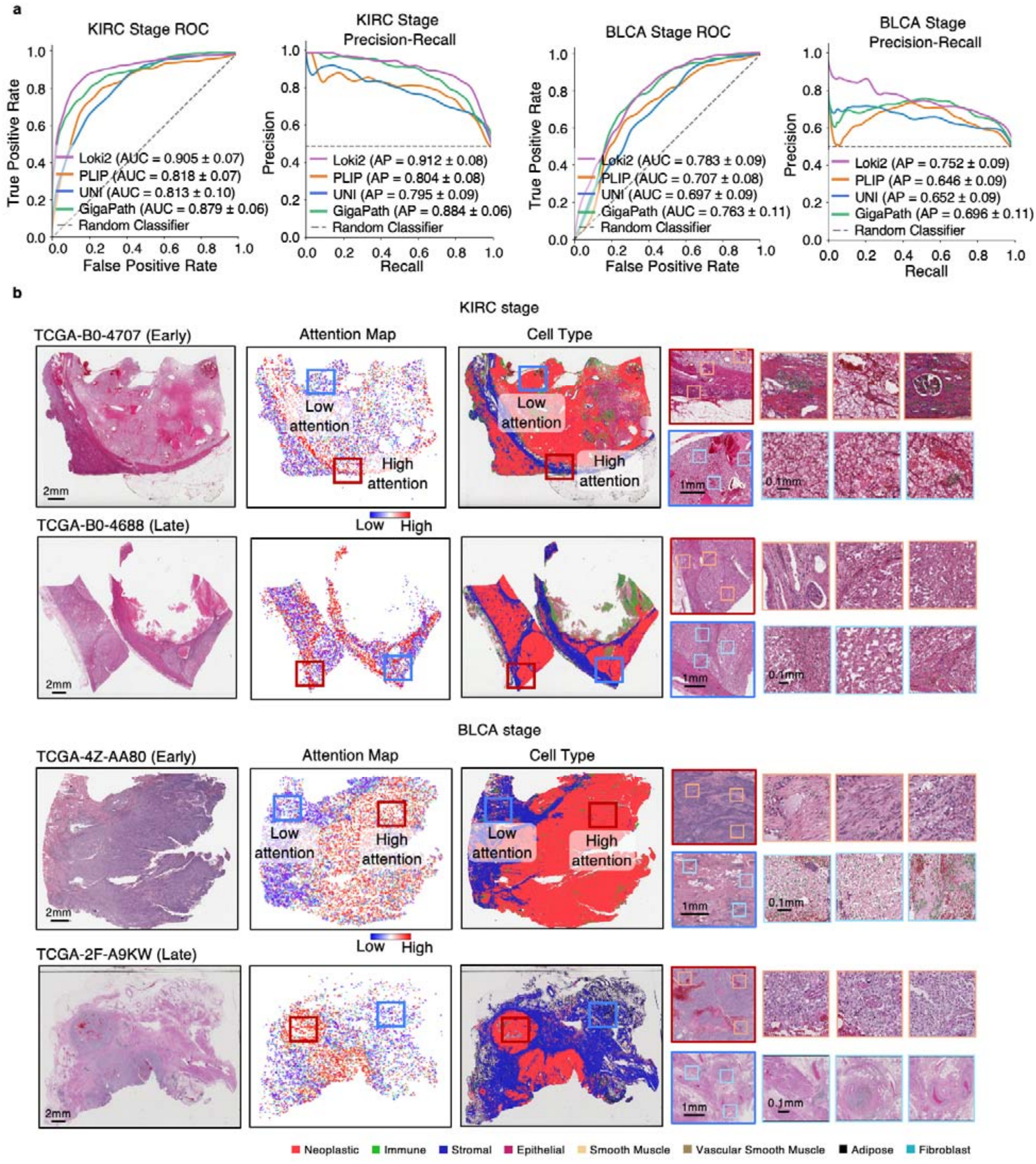
Cell-level multiple instance learning. **a**, Five-fold cross-validation ROC and Precision-Recall curves for KIRC and BLCA stage prediction using MIL model (Loki2) trained on identical single-cell morphology embeddings. Four WSIs foundation models are compared: Loki2 (our model, purple), PLIP (orange), UNI (blue), and GigaPath (green). Left panels show ROC curves and right panels show Precision-Recall curves. **b**, Representative visualizations for KIRC (top two rows) and BLCA (bottom two rows) across Early- and Late-stage samples. Columns from left to right are original H&E whole-slide image, single-cell attention heatmap (red: high attention, blue: low attention), Loki2-decoded cell type map; and multi-resolution ROIs corresponding to high-attention (red box) and low-attention (blue box) regions. Neoplastic cells (red) predominantly drive high attention scores in both cancers.

### Morphology-associated molecular features nominate clinically relevant biomarkers

We next asked whether molecular features retrieved from single-cell morphology could identify candidate biomarkers associated with tumor grade and patient outcomes. We applied the trained Loki2 model to an external cohort from the Prostate, Lung, Colorectal and Ovarian (PLCO) Cancer Screening Trial^44^, comprising whole-slide H&E images from 100 colorectal cancer patients (50 low-grade and 50 high-grade; **Fig. 7a**). Without retraining or cohort-specific adaptation, we performed single-cell morphology-to-transcriptome retrieval in this cohort by matching Loki2-derived cellular morphology embeddings to an independent scRNA-seq reference atlas^45^. This enabled reconstruction of the tumor microenvironment directly from histology alone. Consistent with previous analysis, the retrieved cellular states showed clear spatial concordance with tissue architecture, with tumor cells predominantly localized to histologically defined tumor regions, supporting the biological validity of the retrieval task (**Fig. 7b**). We also observed a higher proportion of tumor cells in high-grade cases (**Fig. 7c**). Using the retrieved tumor-cell transcriptomic profiles, we then performed differential expression analysis between low-grade and high-grade cases. We identified a three-gene signature using the top three significantly differentially expressed genes, S100A6, S100A11, and KRT8, that was consistently elevated in high-grade tumors (**Fig. 7d**). To evaluate clinical relevance independently, we next tested this three-gene signature in an independent TCGA colorectal cancer cohort (n = 151), derived from the rectal adenocarcinoma dataset. Patients were stratified into high- and low-risk groups using a composite score derived from S100A6, S100A11, and KRT8 expression (p value = 0.0089, **Fig. 7e**). This combined signature significantly separated overall survival outcomes. Together, these findings connect morphology-associated molecular recovery to patient-level relevance by showing that transcriptomic retrieval can nominate candidate biomarkers for evaluation of clinical associations using independently measured molecular data.

**Fig 7:**
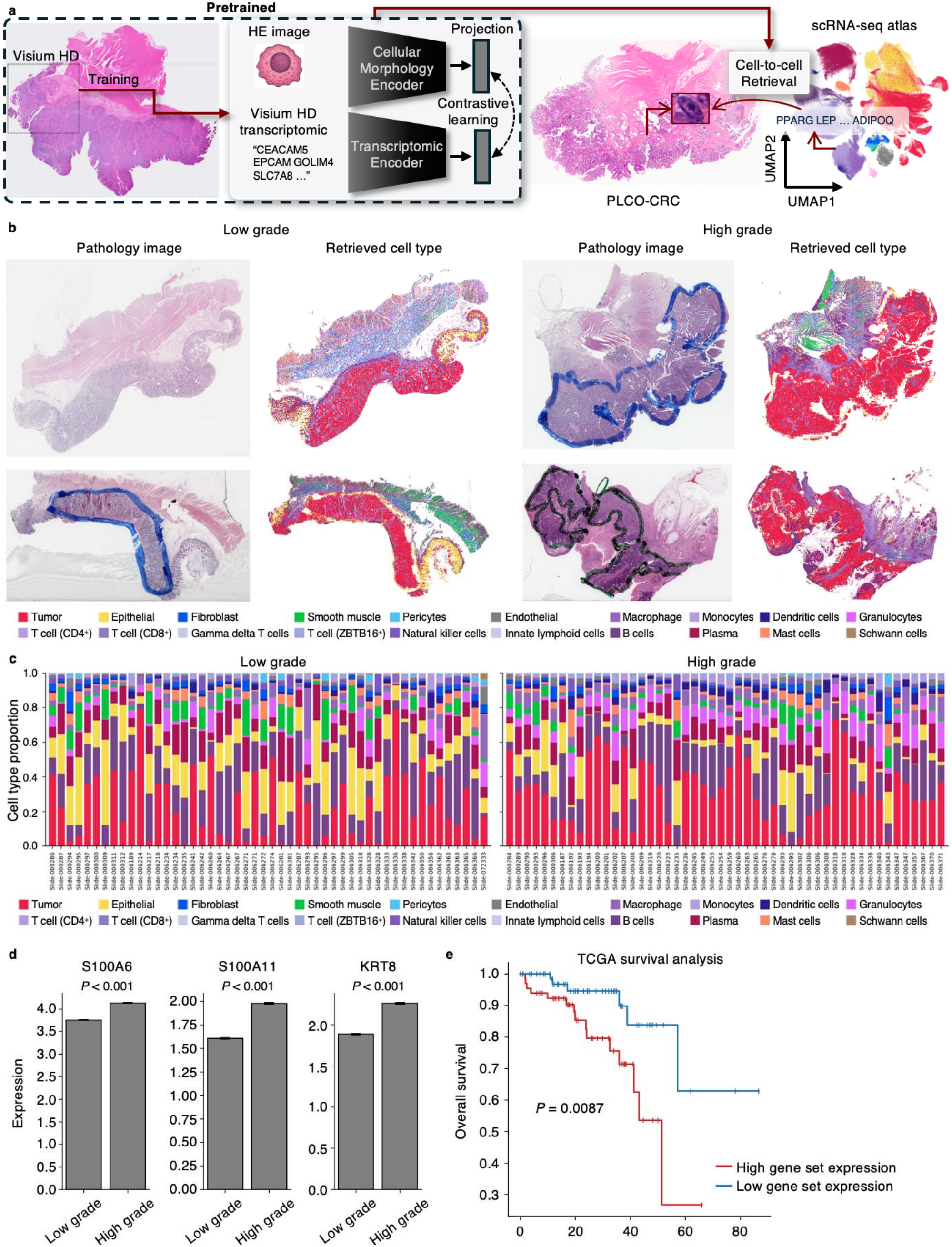
Morphology-to-transcriptome retrieval at single-cell resolution on PLCO cohort. **a**, Schematic overview of single-cell morphology-to-transcriptome retrieval on PLCO cohort using Loki2. **b**, Two PLCO low-grade CRC examples and two PLCO high-grade CRC examples illustrating Loki2 cell segmentation and annotation alongside histology images. **c**, Loki2 predicted cell-type proportions for low-grade and high-grade CRC samples. **d**, Expression levels of top 3 differently expressed genes (S100A6, S100A11, and KRT8) in low-grade versus high-grade tumors. Statistical significance was assessed using a two-sided Wilcoxon rank-sum test. **e**, Kaplan–Meier survival analysis stratified by gene set (S100A6, S100A11, and KRT8) expression levels on TCGA colorectal cancer patients. Statistical significance was assessed using log-rank test.

## Discussion

A major contribution of this work is the introduction of UniSeg, a large-scale multimodal resource that unifies histopathology, cell-level segmentation, lineage annotations, and spatially resolved transcriptomics. By harmonizing 15 million cells across 15 tissues and integrating high-resolution molecular measurements with histology-inferred profiles, UniSeg addresses the data scarcity, tissue bias, and modality fragmentation that have long constrained progress in computational pathology. Much as ImageNet^46^ and LAION^47^ catalyzed breakthroughs in computer vision and vision–language modeling, UniSeg provides the multimodal substrate necessary for next-generation pathology and spatial-omics foundation models, particularly those seeking to anchor morphological abstraction in molecular reality. Importantly, OmiCLIP serves only as one source of molecular supervision during training and does not define the Loki2 model itself.

Beyond dataset scale, this work marks a conceptual shift in how AI systems engage with computational biology: Loki2 treats the single cell but not the patch or tile as the fundamental computational unit. Clinically significant processes, from immune exclusion to epithelial-mesenchymal transition, manifest as continuous shifts in cell states that are frequently obscured by coarse patch-based averaging. By producing high-dimensional morphological embeddings for millions of nuclei, Loki2 aligns computational pathology with the true resolution at which tissues develop, respond to injury, and progress toward malignancy. This single-cell framing enables fine-grained modeling of microenvironmental interactions, transitional cell states, and evolutionary trajectories, offering sharper biological insight into localized or rare cellular patterns that disproportionately influence clinical outcomes.

A central innovation of this study is the demonstration that morphological phenotypes and molecular states are deeply and systematically coupled, and that this relationship can be learned directly from routine H&E images. Through cross-modal contrastive pretraining, Loki2 establishes a shared latent manifold in which visually similar cells map to transcriptionally similar profiles. These embeddings transform the H&E slide into a quantitative, molecularly interpretable data source. By grounding nuclear morphology within a gene expression coordinate system, Loki2 provides a principled and scalable bridge between histopathology and genomics, enabling the extraction of molecular biomarkers in settings where sequencing is limited, cost-prohibitive, or historically unavailable. These embeddings further enable the prediction of multiplexed protein patterns from morphology alone, allowing high-fidelity *in silico* immunostaining in the absence of specialized imaging. Such virtual staining expands the molecular reach of H&E and provides a scalable alternative when laboratory assays are constrained.

Leveraging this cross-modal alignment, Loki2 enables zero-shot *in silico* spatial transcriptomics, reconstructing tissue-wide molecular landscapes by retrieving transcriptionally matched neighbors from scRNA-seq reference atlases. Importantly, this retrieval-based strategy offers a key advantage over regression-based approaches: because each inferred expression vector corresponds to an experimentally measured transcriptome, the reconstruction preserves native gene-gene covariance and maintains biologically realistic expression geometry. This capability democratizes access to spatial molecular information, allowing researchers and clinicians to perform virtual sequencing on standard H&E slides across biobanks, retrospective cohorts, and clinical trials where spatial transcriptomics is infeasible or unavailable.

One of the key limitations of patient-level prediction in Loki2 is its dependence on sufficiently large and well-annotated cohorts. Early metastasis is a representative example: although histopathology-based modeling can capture morphology, tumor–stroma interactions, and immune context associated with metastatic potential, metastasis is a dynamic and multi-stage process that cannot be fully represented by static tissue sections from limited regions. Loki2 may help address part of this challenge by extracting single-cell morphology features and linking them to latent molecular states. However, the limited availability of large, standardized datasets with matched primary and metastatic samples still constrains model training, evaluation, and generalization. Thus, integrating histopathology with longitudinal clinical data and multimodal molecular profiling will be important for more comprehensive disease condition such as metastatic progression.

As a single-cell pathology foundation model, Loki2 provides a general, biologically interpretable representation for investigating morphology–molecular relationships. The present study evaluates the cell-level interpretability, molecular recoverability and clinical relevance of this representation, but does not include perturbation experiments to establish causality. An important future direction is to combine Loki2 with targeted genetic or pharmacological perturbations and paired morphological and molecular measurements to determine how specific regulatory programs shape cellular morphology and whether these relationships persist under intervention.

Extending interpretability across spatial scales is another important future direction. Our current analysis focuses on nuclear morphology at the single-cell level, the fundamental unit of representation in Loki2. Tissue-level phenotypes, such as necrosis, immune infiltration, stromal activation, epithelial differentiation and tumor grade, also depend on the composition and spatial organization of multicellular regions. Integrating annotations of spatially defined tissue regions/units with explicit aggregation of cellular representations could link interpretable nuclear features to these broader tissue phenotypes, clarifying how cellular states and their spatial relationships contribute to pathological architecture.

In Loki2, retrieval-based molecular inference depends on the availability and biological compatibility of an appropriate scRNA-seq reference. When relevant cell types or disease states are absent, retrieval cannot recover their missing profiles and may instead assign cells to biologically mismatched reference populations. If no suitable reference is available, we recommend generating scRNA-seq data from the tissue and condition of interest to capture the relevant cellular diversity. This limitation applies specifically to transcriptomic retrieval, rather than to Loki2’s morphological representations or applications that do not require an scRNA-seq reference.

## Acknowledgments

This work was supported in part by the National Institute of Health (NIH) R35GM150460 (G.W.), R01HL179815 (G.W.), R01AG089925 (Q.M.), R01GM152585 (Q.M.), R01DK138504 (Q.M.), P01AI177687 (Q.M.), P01CA278732 (Q.M.), and U54AG075931 (Q.M.). This work was also supported by the Pelotonia Institute of Immuno-Oncology (PIIO) at The Ohio State University. The content is solely the responsibility of the authors and does not necessarily represent the official views of the NIH and the PIIO.

## Author Contributions

These authors contributed equally: Weiqing Chen, Qing Li, and Hao Cheng.

G.W., Q.M., and S.L. supervised the study. W.C., G.W., Q.M., and S.L. designed and developed the Loki2 model and platform. W.C., Q. L., H.C., P.Z., T.T., C.L., and Y.Y. analyzed the data.

G.W., Q.M., W.C., Q.L., and P.Z. wrote the manuscript. A.M. and M.R. provided in-house lung Xenium data. S.A. and S.H.L. provided in-house esophageal patient data. Y.C., X.W., Y.X., X.Z., W.W., and A.M., were involved in the discussion and helped improve the manuscript. All authors approved the final version of the manuscript.

## Ethics declarations

Competing interests

The authors declare no competing interests.

## Figure Legend

**Fig. 1: Overview of the study**.

**a**, The workflow of UniSeg resource preparation.

**b**, The left diagram illustrates the size of UniSeg resource in different organs. The right diagram shows the example image patches with cell boundary, type annotation, and paired marker gene expressions from UniSeg.

**c**, The workflow of Loki2 model training.

**d**, Overview of the Loki2 framework.

**Fig. 2: Benchmarking Loki2 for single-cell segmentation, typing, and generalization. a**, Box plots comparing mPQ scores for Loki2, CellViT, and HoVer-Net across 9,573 test instances from 15 tissue types (sample sizes: colon 2,552; lung 1,300; heart 1,253; prostate 1,000; kidney 753; breast 500; brain 500; skin 400; lymph node 389; liver 250; ovarian 200; small intestine 150; pancreatic 150; bladder 126; cervix 50). In the box plots, the middle line represents the median, the box boundaries indicate the interquartile range, and the whiskers extend to data points within 1.5 times the interquartile range.

**b**, Representative examples comparing segmentation and cell-type annotation results from Loki2, CellViT, HoVer-Net, and BiomedParse with ground truth.

**c**, Box plots of weighted Dice and binary Dice scores for Loki2 and BiomedParse on the same test instances. Box plot elements follow the conventions described in (a).

**d**, Four TCGA examples illustrating Loki2 cell segmentation and annotation alongside pathologist-annotated tumor regions.

**e**, SHAP analysis of the seven morphology features predicted from H&E images of 20 held-out samples. Left, mean |SHAP| value for each of seven morphology features, with stacked segments showing the contributing PCs. Middle and right, SHAP beeswarm plots for chromatin density and nuclear enlargement. Each point is one cell; color shows the z-scored PC value.

**f**, Representative DAPI nuclei with low and high predicted chromatin density and nuclear enlargement.

**Fig. 3: Morphology-based pseudotime inference**.

**a**, H&E images, Loki2 predicted cell types, cellular morphology embeddings, and inferred morphological pseudotime for three CRC samples. For cellular morphology embeddings, the samples were visualized by mapping their three UMAP components, that computed from cellular morphology embeddings to the red, green, and blue channels, respectively.

**b**, UMAP visualization of cellular morphology embeddings from the three CRC samples, colored by inferred morphological pseudotime and somatic mutations from the original study, respectively.

**c**, H&E image, Loki2 identified epithelial and neoplastic cells, cellular morphology embeddings, and inferred morphological pseudotime for the prostate cancer sample. For cellular morphology embeddings, the samples were visualized by mapping their three UMAP components, that computed from cellular morphology embeddings, to the red, green, and blue channels, respectively.

**d**, UMAP visualization of cellular morphology embeddings from the prostate cancer sample, colored by inferred morphological pseudotime and region types from the original study, respectively.

**Fig. 4: Morphology-to-transcriptome retrieval at single-cell resolution**.

**a**, Schematic overview of single-cell morphology-to-transcriptome retrieval using Loki2.

**b**, Left panels show cell-type annotations from the pathologist and from Loki2, including corresponding zoom-in regions. Right panels show example gene-expression maps from Visium HD (green dashed box region) alongside predictions from Loki2, GHIST, and iStar, with SSIM values computed relative to the Visium HD ground truth.

**c**, Visualization of samples by mapping the top three PCA components that fit on Visium HD gene expression and then applied to gene expression predictions from Loki2, GHIST, and iStar, to the red, green, and blue channels, respectively.

**d**, Gene expression prediction performance comparison using Spearman correlation and Pearson correlation of highly expressed gene ranks predicted by Loki2, GHIST, and iStar, respectively. Statistical significance was assessed using a one-sided Mann–Whitney U test; *P-value < 5×10^-2^, **P-value < 1×10^-2^, ***P-value < 1×10^-3^, and ****P-value <1×10^-4^. In the box plots, the middle line represents the median, the box boundaries indicate the interquartile range, and the whiskers extend to data points within 1.5 times the interquartile range.

**Fig. 5: *In silico* immunostaining from standard histology**.

**a**, The workflow of Loki2 *in silico* immunostaining from standard histology.

**b**, Benchmarking results showing MAE across four CyCIF markers, including CD45RO, Ki67, CD4, and CD45. Statistical significance was assessed using a one-sided Mann–Whitney U test; *P-value < □ 5×10^-2^, **P-value < □ 1×10^-2^, ***P-value < □ 1×10^-3^, and ****P-value <1×10^-4^.

**c**, Benchmark results showing SSIM values for the same four CyCIF markers.

**d**, An example CRC H&E image patch.

**e**, Visualization of raw H&E, virtual stain images generated by Loki2 and ROSIE, respectively, across CD45RO, PanCK, and CD45.

**f**, Benchmarking results showing MAE for PanCK in both IHC and CyCIF. Statistical significance was assessed using a one-sided Mann–Whitney U test; *P-value < □ 5×10^-2^, **P-value < □ 1×10^-2^, ***P-value < □ 1×10^-3^, and ****P-value <1×10^-4^.

**g**, Benchmarking results showing SSIM values for PanCK in IHC and CyCIF.

**h**, An example colorectal cancer H&E image patch.

**i**, Visualization of the raw H&E image and virtual stain images generated by Loki2 and ROSIE, respectively.

**j**, Example whole-slide H&E images and corresponding Loki2-predicted PanCK protein intensity with zoom-in regions from PLCO (left) and TCGA (right) cohorts.

**k**, Quantitative comparison of Z-score of Loki2 predicted PanCK intensity inside and outside pathologist annotated tumor regions across multiple samples from PLCO and TCGA cohorts. In the box plots, the middle line represents the median, the box boundaries indicate the interquartile range, and the whiskers extend to data points within 1.5 times the interquartile range.

**Fig. 6: Cell-level multiple instance learning**.

**a**, Five-fold cross-validation ROC and Precision-Recall curves for KIRC and BLCA stage prediction using MIL model (Loki2) trained on identical single-cell morphology embeddings. Four WSIs foundation models are compared: Loki2 (our model, purple), PLIP (orange), UNI (blue), and GigaPath (green). Left panels show ROC curves and right panels show Precision-Recall curves.

**b**, Representative visualizations for KIRC (top two rows) and BLCA (bottom two rows) across Early- and Late-stage samples. Columns from left to right are original H&E whole-slide image (WSI), single-cell attention heatmap (red: high attention, blue: low attention), Loki2-decoded cell type map; and multi-resolution ROIs corresponding to high-attention (red box) and low-attention (blue box) regions. Neoplastic cells (red) predominantly drive high attention scores in both cancers.

**Fig. 7: Morphology-to-transcriptome retrieval at single-cell resolution on PLCO cohort. a**, Schematic overview of single-cell morphology-to-transcriptome retrieval on PLCO cohort using Loki2.

**b**, Two PLCO low-grade CRC examples and two PLCO high-grade CRC examples illustrating Loki2 cell segmentation and annotation alongside histology images.

**c**, Loki2 predicted cell-type proportions for low-grade and high-grade CRC samples.

**d**, Expression levels of top 3 differently expressed genes (S100A6, S100A11, and KRT8) in low-grade versus high-grade tumors. Statistical significance was assessed using a two-sided Wilcoxon rank-sum test.

**e**, Kaplan–Meier survival analysis stratified by gene set (S100A6, S100A11, and KRT8) expression levels on TCGA colorectal patients. Statistical significance was assessed using log-rank test.

**Extended Data Fig. 1: Example of OmiCLIP transcriptomic embeddings**.

The OmiCLIP transcriptomic embeddings extracted from 4 µm bins Visium HD data of a CRC sample. The sample is visualized by mapping its three UMAP components that computed from cell transcriptomics embeddings to the red, green, and blue channels, respectively.

**Extended Data Fig. 2: Example cell embeddings from UniSeg**.

H&E images, clusters of cell morphology embeddings, clusters of cell transcriptomic embeddings, and clusters of combined morphology–transcriptomic embeddings from five representative samples: normal liver, skin cancer, colorectal cancer, breast cancer, and lung cancer tissues.

**Extended Data Fig. 3: Benchmarking Loki2 for single-cell segmentation, typing, and generalization**.

**a**, Box plots comparing mF1 scores for Loki2, CellViT (remapped labels), and HoVer-Net (remapped labels) across 9,573 test instances from 15 tissue types (sample sizes: colon 2,552; lung 1,300; heart 1,253; prostate 1,000; kidney 753; breast 500; brain 500; skin 400; lymph node 389; liver 250; ovarian 200; small intestine 150; pancreatic 150; bladder 126; cervix 50). In the box plots, the middle line represents the median, the box boundaries indicate the interquartile range, and the whiskers extend to data points within 1.5 times the interquartile range.

**b**, Representative examples comparing segmentation and cell-type annotation results from Loki2, CellViT, HoVer-Net, and BiomedParse with ground truth.

**c**, Box plots of pixel accuracy for Loki2 and BiomedParse on the same test instances. Box plot elements follow the conventions described in (a).

**d**, Four TCGA examples illustrating Loki2 cell segmentation and annotation alongside pathologist-annotated tumor regions.

**e**, Loki2 predicted cell-type proportions for TCGA BRCA HER2^+^ samples.

**Extended Data Fig. 4: Cell type annotation out-of-distribution benchmark. a**, WSIs of the held-out samples.

**b**, Box plots comparing mPQ scores for Loki2, CellViT (remapped labels), and HoVer-Net (remapped labels) across 7 held-out whole-slide images from distinct patients (n=50 for each sample). In the box plots, the middle line represents the median, the box boundaries indicate the interquartile range, and the whiskers extend to data points within 1.5 times the interquartile range.

**c**, Box plots comparing mPQ scores for Loki2, CellViT (remapped labels), and HoVer-Net (remapped labels) across 8 held-out patients’ whole-slide images (n=50 for each patient). In the box plots, the middle line represents the median, the box boundaries indicate the interquartile range, and the whiskers extend to data points within 1.5 times the interquartile range.

**Extended Data Fig. 5: Histological structure and morphology-based similarity tree**.

**a**, Representative pathology image (left) and corresponding region type annotation (right) of a dorsolateral prefrontal cortex tissue section. Black dots represent cells detected by Loki2.

**b**, Hierarchical clustering of region types based on their Loki2 cell feature similarity.

**Extended Data Fig. 6: Cell type classification performance comparison across models**. Bar plot comparing weighted F1 score and accuracy for cell type classification using Loki2 and different pathology foundation models as backbone, including PLIP, GigaPath and UNI.

**Extended Data Fig. 7: Interpretable nuclear-morphology analysis of the Loki2 embeddings**.

**a**, Study design: 18 esophageal patient samples with paired H&E and DAPI staining. **b**, SHAP analysis of the seven morphology features predicted from H&E images of 18 independent esophageal patient samples. Left, mean |SHAP| value for each of seven morphology features, with stacked segments showing the contributing PCs. Middle and right, SHAP beeswarm plots for chromatin density and nuclear enlargement. Each point is one cell; color shows the z-scored PC value.

**c**, Cumulative distributions of each morphology feature in the top 25% group (red) versus bottom 25% group (blue), for held-out H&E images, esophageal H&E images, and esophageal DAPI images. P values, two-sided Mann–Whitney U test. Eosinophilic foci cannot be scored in DAPI.

**d**, Bar plot comparing weighted F1 score and accuracy for cell type classification using Loki2 and 7 conventional morphological features respectively.

**Extended Data Fig. 8: H&E and DAPI images of 18 esophageal cancer specimens**.

Paired H&E and DAPI whole-slide images acquired from the 18 patients.

**Extended Data Fig. 9: Diffusion map of cell morphology embeddings across different tissues**.

Two-dimensional diffusion map projections (diffusion component 2 vs. diffusion component 3) of single-cell morphology embeddings are shown for **a**, skin, brain, and **b**, two CRC patient samples, where each point represents an individual cell.

**Extended Data Fig. 10: Cross-tissue single-cell morphology–transcriptome similarity analysis**.

**a**, Spatial visualization of cell embeddings derived from morphology (top rows) and transcriptomics (bottom rows) across multiple tissues. Each point represents a cell, colored by its low-dimensional embedding representation.

**b**, Cross-tissue comparison of embedding similarity. Boxplots showing the distribution of cosine similarity scores across different tissues.

**Extended Data Fig. 11: Root cell selection of morphology-based pseudotime inference. a**, Selected root cells for pseudotime initialization in the three CRC samples.

**b**, Selected root cell for pseudotime initialization in the prostate cancer sample.

**Extended Data Fig. 12: Morphological pseudotime correlated gene analysis**.

**a**, Scatter plots showing expression of representative high correlated genes (TMSB4X and IGFBP2) and low correlated genes (CTSB) along inferred pseudotime.

**b**, Heatmap showing the enrichment significance of hallmark pathways across three gene groups stratified by correlation levels. Columns represent pathways, and rows correspond to gene groups. Color intensity indicates the relative p-value within each row (row-wise normalization from minimum to maximum), with darker colors representing stronger enrichment.

**Extended Data Fig. 13: Workflow of morphology-to-transcriptome retrieval at single-cell resolution**.

**a**, Schematic illustration of the morphology-to-transcriptome retrieval with contrastive learning. **b**, UMAP visualization of morphology embeddings and transcriptomic embeddings, colored by mapping its three UMAP components that were computed from cell morphology embeddings and from cell transcriptomics embeddings to the red, green, and blue channels, respectively. **c**, UMAP of morphology embeddings and transcriptomics embeddings colored by morphology modality and transcriptomics modality, respectively.

**d**, Visualization of the sample by mapping its three UMAP components that computed from cell morphology embeddings and from cell transcriptomics embeddings to the red, green, and blue channels, respectively.

**e**, Schematic illustration of morphology-to-transcriptome retrieval applied to Visium HD data.

**Extended Data Fig. 14: Marker gene validation of cell type annotations in paired and external scRNA-seq datasets**.

Dot plots showing the expression of canonical marker genes across annotated cell types in **a**, paired scRNA-seq data and **b**, an independent external scRNA-seq dataset. Rows correspond to cell types, and columns represent selected marker genes grouped by lineage. Dot size indicates the fraction of cells within each cell type expressing the gene, and color intensity represents the mean expression level.

**Extended Data Fig. 15: Examples of gene expression prediction**.

Left panels show cell-type annotations from the pathologist and from Loki2, including corresponding zoom-in regions. Right panels show example gene-expression maps from Visium HD (green dashed box region) alongside predictions from Loki2, GHIST, and iStar, with SSIM values computed relative to the Visium HD ground truth.

**Extended Data Fig. 16: Comparison of gene expression prediction performance**.

SSIM values for the top 100 expressed genes across four test samples, evaluated using Loki2, GHIST, and iStar, respectively. Statistical significance was assessed using a one-sided Mann– Whitney U test; *P-value <□ 5×10^-2^, **P-value □ < 1×10^-2^, ***P-value □ < 1×10^-3^, and ****P-value

<1×10^-4^. In the box plots, the middle line represents the median, the box boundaries indicate the interquartile range, and the whiskers extend to data points within 1.5 times the interquartile range.

**Extended Data Fig. 17: PCA visualization of gene expression prediction**.

Visualization of samples by mapping the top three PCA components that fit on Visium HD gene expression and then applying them to gene expression predictions from Loki2, GHIST, and iStar, to the red, green, and blue channels, respectively.

**Extended Data Fig. 18: Morphology-to-transcriptome retrieval at single-cell resolution finetuned on Xenium data**.

**a**, Schematic overview of single-cell morphology-to-transcriptome retrieval using Loki2.

**b**, Left panels show cell-type annotations from the pathologist and from Loki2, including corresponding zoom-in regions. Right panels show example gene-expression maps from Visium HD ground truth (green dashed box region) alongside predictions from Loki2, with SSIM values computed relative to the ground truth.

**c**, Gene expression prediction performance comparison using Spearman correlation and Pearson correlation of highly expressed gene ranks, and SSIM values for the top 100 expressed genes predicted by Loki2 (finetuned on Visium HD data), Loki2 (finetuned on Xenium data), GHIST, and iStar, respectively. In the box plots, the middle line represents the median, the box boundaries indicate the interquartile range, and the whiskers extend to data points within 1.5 times the interquartile range.

**Extended Data Fig. 19: Morphology-to-transcriptome retrieval at single-cell resolution finetuned on Xenium data**.

Left panels show cell-type annotations from the pathologist and from Loki2, including corresponding zoom-in regions. Right panels show example gene-expression maps from Visium HD ground truth (green dashed box region) alongside predictions from Loki2, with SSIM values computed relative to the ground truth.

**Extended Data Fig. 20: Morphology-to-transcriptome retrieval at single-cell resolution using external scRNA-seq reference data**.

**a**, Left panels show cell-type annotations from the pathologist and from Loki2, including corresponding zoom-in regions. Right panels show example gene-expression maps from Visium HD ground truth (green dashed box region) alongside predictions from Loki2 using external scRNA-seq reference data, with SSIM values computed relative to the ground truth.

**b**, SSIM values for the top 100 expressed genes retrieved by Loki2 from external reference scRNA-seq data under different mixing ratios with paired reference scRNA-seq data (paired data only; paired 80% + external 20%; paired 60% + external 40%; paired 20% + external 80%; external data only). In the box plots, the center line represents the median, the box bounds indicate the interquartile range, and whiskers extend to 1.5× the interquartile range.

**Extended Data Fig. 21: Morphology-to-transcriptome retrieval at single-cell resolution using external scRNA-seq reference data**.

Left panels show cell-type annotations from the pathologist and from Loki2, including corresponding zoom-in regions. Right panels show example gene-expression maps from Visium HD ground truth (green dashed box region) alongside predictions from Loki2 using external scRNA-seq reference data, with SSIM values computed relative to the ground truth.

**Extended Data Fig. 22: Morphology-molecular coupling analysis**.

**a**, Genes ranked by mutual information (MI) with morphology embeddings, split into five equal groups from G1 (highest) to G5 (lowest).

**b**, Scatter plots of expression versus top PC score

**c**, Stacked bars show the proportions of linear probing genes ranked by Spearman correlation across G1–G5.

**d**, Representative genes, summarized functional annotations, mean number of expressing cell types and mean cell type specificity for the five gene groups (G1–G5).

**Extended Data Fig. 23: Gene expression versus morphology-derived principal component scores for genes with high and low mutual information**.

Scatter plots of high MI gene (top) expression versus top PC score and low MI gene (bottom) expression versus top PC score from colorectal cancer sample and lung cancer sample respectively.

**Extended Data Fig. 24: *In silico* immunostaining from standard histology**.

**a**, Benchmarking results showing MAE and SSIM for CD45RO, Ki67, CD4, and CD45 CyCIF markers across linear regression (LR), support vector regression (SVR), multilayer perceptron (MLP), LightGBM, and ROISE.

**b**, Visualization of raw H&E images and corresponding virtual stain images generated by Loki2 and ROSIE, respectively, for Ki67 and CD4.

**c**, Benchmarking results showing MAE and SSIM for PanCK (IHC and CyCIF) and CD3 (IHC) markers across LR, SVR, MLP, LightGBM, and ROISE.

**d**, Example whole-slide H&E images and corresponding Loki2-predicted PanCK protein intensity with zoom-in regions from PLCO and TCGA cohorts.

**Extended Data Fig. 25: Performance of cell-level MIL in COAD stage prediction**. Performance evaluation of staging prediction for COAD. Left panel shows the ROC curves, and right panel shows the Precision-Recall curves for the four foundation models: Loki2, PLIP, UNI, and GigaPath. AUC values are reported across multiple 5-fold cross-validation folds. The dashed gray line represents the performance of a random classifier.

**Extended Data Fig. 26: Example cell-level MIL attention heatmaps**.

Additional representative cases for KIRC early- and late-stage samples. Columns (left to right): original H&E whole-slide image, single-cell attention heatmap, Loki2-decoded cell type map, and high-resolution ROIs corresponding to high (red box) and low (blue box) attention scores with detailed views.

**Extended Data Fig. 27: Example cell-level MIL attention heatmaps**.

Additional representative cases for BLCA early- and late-stage samples. Columns (left to right): original H&E whole-slide image, single-cell attention heatmap, Loki2-decoded cell type map, and high-resolution ROIs corresponding to high (red box) and low (blue box) attention scores with detailed views.

**Extended Data Fig. 28: Example cell-level MIL attention heatmaps**.

Representative cases comparing COAD early- and late-stage samples. Columns (left to right): original H&E whole-slide image, single-cell attention heatmap, Loki2-decoded cell type map, and high-resolution ROIs corresponding to high (red box) and low (blue box) attention scores with detailed views.

## Supplementary information

Supplementary Table 1-4, and Supplementary Note 1.

## Methods

### UniSeg Dataset curation

To build a unified multimodal resource for training a foundation model capable of resolving cellular states from H&E images, we curated UniSeg, a large-scale multimodal resource comprising approximately 15 million cells paired with expert-validated nuclear masks, cell-type annotations, and matched spatial transcriptomic profiles. We collected the H&E images with paired spatial transcriptome profiles from publicly accessible sources, including technologies of 10x Visium, Visium HD, and Xenium, aggregated from 65 studies spanning 15 tissue types (Supplementary Table 1, 2). We first detect the cell boundaries from the H&E images using automated deep learning-based segmentation and subsequently refined by human experts (Supplementary Note 1). To obtain paired spatial transcriptomes for each detected cell, for high-resolution platforms including Visium HD and Xenium, transcript abundance vectors were directly extracted at cellular resolution. For spot-level Visium data, we applied Thor^13^ to generate histology-inferred single-cell profiles. For each curated cell with a paired H&E image, cell boundary, and spatial transcriptome profiles, we extracted its morphology embeddings from the H&E image using OmiCLIP^11^ image encoder, and its transcriptomic embeddings from spatial transcriptomic profiles using OmiCLIP transcriptomic encoder. These resulting two modality embeddings were concatenated to generate unified multimodal representations, which were subsequently used to obtain initial cell-type labels with ChatGPT-assisted annotation^48^, followed by expert review and correction (Supplementary Note 1). We manually reviewed all AI-generated annotations, comparing them against cell-type labels reported in the original publications for 111 samples, while for the remaining 102 samples without reported labels, annotations were confirmed based on morphological assessment through consultation with pathologists. Together, this workflow yielded a harmonized dataset encompassing 22 major cellular lineages, serving as the foundation for training the Loki2 model.

### Loki2 model training

Loki2 consisted of a U-Net-shaped^15^ encoder–decoder network augmented with a ViT^16^ backbone to jointly model cell morphology, segmentation, and transcriptomic variation. The convolutional encoder performed cell-level feature extraction and segmentation, while the integrated ViT module encodes high-dimensional morphology embeddings from image patches. For segmentation, Loki2 employed a Dice loss:

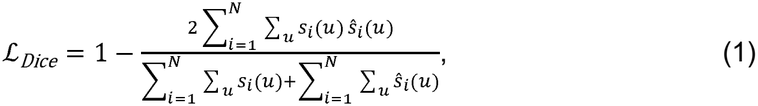

where *s*_*i*_(*u*) and *ŝ*_*i*_(*u*) denote the ground-truth and predicted binary masks for pixel *u* of *i* cell . Parallel to this, transcriptomic embeddings for the same cells were obtained using the OmiCLIP^11^ encoder. Loki2 employed a contrastive learning objective that aligns the morphology embeddings with their paired transcriptomic embeddings and reconstructs the transcriptomic embeddings. The contrastive loss is calculated as:

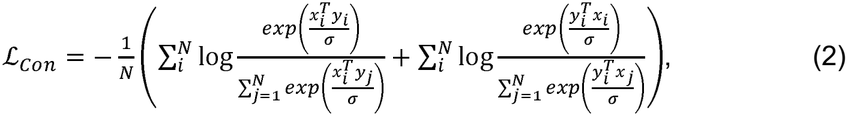

where *x*_*i*_ denotes *y*_*j*_ the normalized morphology and transcriptomic embeddings, respectively. *N* denotes the batch size, while represents the temperature parameter, which *σ* scales the similarity scores and controls the sharpness of this separation. This contrastive loss drives the alignment between morphology and transcriptomic representations by bringing matched pairs closer and pushing unmatched pairs apart in the shared embedding space. Two task-specific decoders operated on the encoded morphology embeddings: one decoder produced cell-type inference maps, and the other reconstructed transcriptomic embeddings, both at single-cell resolution. The cell-type classification decoder was optimized using a binary cross-entropy (BCE) loss:

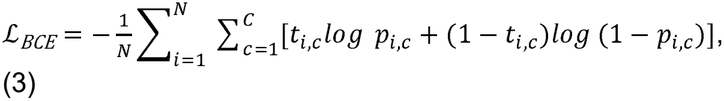

where t_i,c_ is the ground-truth label indicating whether cell i belongs to class *c*, and *p*_*i,c*_ is the predicted probability for that class. The transcriptomic reconstruction decoder was trained using a mean squared error (MSE) loss between the predicted and ground-truth transcriptomic embeddings:

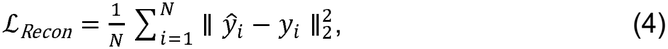

where *y*_*i*_ denotes the ground-truth transcriptomic embedding for cell *i, ŷ*_*i*_, represents the H&E patch was standardized to a size of 256 × 256 pixels, and *N* the dataset was split into train:validation:test partitions of 1:1:1. The model was pretrained for 50 epochs, using eight NVIDIA H200 141-GB GPUs with a local batch size of 16. The output vectors with dimensions of 1280 were optimized by minimizing the total loss on a given batch. All experiments were run in Python v.3.10. Detailed software versions are CUDA v.13.0; torch v.2.9.1; torchvision v.0.24.1; scipy v.1.15.2; scikit-learn v.1.7.2; pandas v.2.3; numpy v.1.26.4; and scanpy v.1.11.5.

### Evaluation of cell segmentation and annotation

We curated the test dataset from the UniSeg database of 9,573 image patches, including 2,552 colon, 1,300 lung, 1,253 heart, 1,000 prostate, 753 kidney, 500 breast, 500 brain, 400 skin, 389 lymph node, 250 liver, 200 ovarian, 150 small intestine, 150 pancreatic, 126 bladder, and 50 cervix tissue images. For CellViT and HoVer-Net, we evaluated we applied the default settings for cell segmentation and annotation. We remapped the fine-grained ground-truth cell types onto their coarse PanNuke parent categories: fibroblasts and cardiac fibroblasts were assigned to stromal, and goblet cells, alveolar pneumocytes, chondrocytes, hepatocytes, pericentral hepatocytes, and proliferating and spinous keratinocytes were assigned to epithelial. Ground truth types with no PanNuke counterpart: smooth muscle, vascular smooth muscle, adipose, cardiomyocyte, endothelial, neuron, glia, oligodendrocyte, and astrocyte, were retained in their original form. We further benchmarked Loki2 against alternative pathology foundation models, including PLIP, GigaPath, and UNI, in the cell type annotation task. For the base line models, we used their pretrained backbones respectively following their pipelines, to extract feature representations from individual cell image patches. We also benchmarked on the seven conventional morphological features (Supplementary Table 3), extracted from each nuclear instance. These frozen embeddings were then used as input to a random forest classifier, which was trained to predict cell type labels on the train and test dataset as Loki2.

#### Evaluation metrics

Multi-class panoptic quality (mPQ): To comprehensively evaluate segmentation performance across both instance- and semantic-level accuracy, we adopted the multi-class panoptic quality (mPQ)^19^. For each class *c*, the panoptic quality *PQ*_*c*_ is defined as:

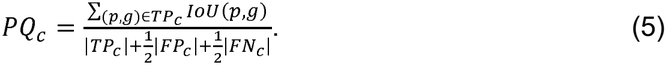

*TP*_*c*_ is the set of true positive (predicted, ground-truth) instance pairs with Intersection-over-Union (IoU) > 0.5, *FP*_*c*_ is false positive predictions (spurious instances), *FN*_*c*_ is false negatives (missed ground-truth instances), and *IoU*(*p, g*) denotes the intersection-over-union between the predicted instance *p* and the ground-truth instance *g*. The mPQ is then obtained by averaging over all classes *C*:

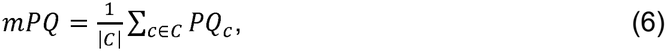

which ranges from 0 to 1.

Multi-class F1 score (mF1): To further assess segmentation performance at the instance level, we computed the multi-class F1 score (mF1), which summarizes the harmonic mean of precision and recall for each class. For each class *c*, true positives *TP*_*c*_ were defined as predicted instances that matched a ground-truth instance with an IoU greater than 0.5, while false positives *FP*_*c*_ and false negatives *FN*_*c*_ were defined as unmatched predictions and unmatched ground-truth instances, respectively:

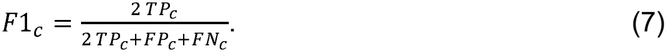

The overall multi-class F1 score is computed by averaging across all classes *C*:

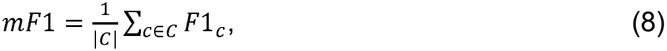

which ranges from 0 to 1.

Binary Dice Score: To quantify overall foreground–background segmentation accuracy, we report the binary dice score, a standard overlap measure widely used in image segmentation. Given a predicted binary mask *P* and ground-truth mask *G*, the Dice score is defined as:

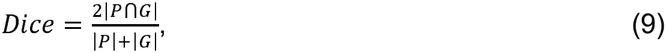

which ranges from 0 to 1.

Weighted Dice: To account for class imbalance, we additionally compute the weighted Dice. For each class *c*, the per-class Dice score *Dice*_*c*_ is computed as above, and a global score is obtained by weighting each class proportionally to its ground-truth pixel frequency *w*_*c*_:

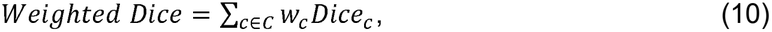

which ranges from 0 to 1.

Pixel accuracy: Pixel accuracy provides a simple measure of semantic segmentation correctness by quantifying the proportion of pixels whose predicted class label matches the ground truth. Formally, pixel accuracy is defined as:

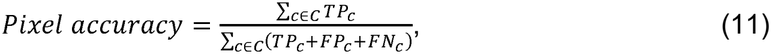

where *TP*_*c*_ denotes correctly classified pixels of class *c*.

Weighted F1 score: To account for class imbalance in cell type annotation, we report the weighted F1 score, where each class-specific F1 score is weighted by its support (i.e., the number of ground-truth instances *N*_*c*_):

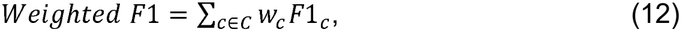

where

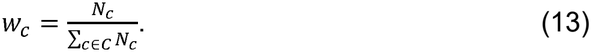

Weighted Accuracy: We also report the weighted accuracy, defined as the average per-class accuracy weighted by the number of ground-truth instances in each class. For each class *c*, the class-wise accuracy is given by *TP*_*c*_/*N*_*c*_, where *TP*_*c*_ denotes the number of correctly predicted instances and *N*_*c*_ is the total number of ground-truth instances for class *c*. The weighted accuracy is then computed as:

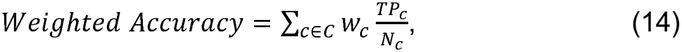

where

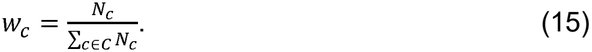

### Interpretable nuclear-morphology analysis of the Loki2 embeddings

Seven conventional morphological features were chosen to span the cytological axes that carry diagnostic weight in routine reporting: nuclear enlargement, nuclear elongation, contour irregularity, chromatin density, chromatin coarseness, chromatin pattern, and eosinophilic foci.

Each feature transcribes an established descriptor from the pathology and quantitative-histology literature into a single scalar (see Supplementary Table 3 for details). All seven features derive from the same nuclear segmentation that defines the cells for which Loki2 embeddings are extracted, so each cell contributes one 1,280-dimensional embedding and one seven-vector of measurements. The discovery cohort was drawn from UniSeg and comprises 100 H&E samples from 96 patients, spanning six source cohorts and six tissue types. Samples were assigned to folds at the patient level, so that all samples from one patient fall on the same side of the split, giving 80 training samples (785,414 cells) and 20 patient-held-out samples (195,814 cells).

A principal-component basis was fitted on the training samples alone by exact covariance eigendecomposition, and the minimal prefix reaching 90% of cumulative training variance was retained. Held-out cells were projected onto this frozen basis. The seven features were standardized within each sample. Component scores and the seven features were then z-scored on training-cell statistics only, so that all attributions and effect sizes are in units of the training target standard deviation. One extremely randomized trees regressor (scikit-learn^49^ ExtraTreesRegressor; 96 trees, maximum depth 18, minimum 20 samples per leaf, half of the features considered per split, no bootstrap) was fitted per feature on all component scores, using up to 1,000 cells per training sample drawn with a deterministic sample-name-derived seed (80,000 cells in total). Each model was explained with exact TreeSHAP^50^ (shap.TreeExplainer, tree_path_dependent perturbation, no approximation) on a balanced subsample of 50 cells per sample (1,000 held-out cells), yielding a signed contribution of each of the 267 components to each explained prediction. Within a feature these contributions are exactly additive, their signed sum equaling the model’s prediction minus its training baseline. Component importance is the mean absolute contribution over explained cells, reported both in target standard deviations and as a within-feature share.

To test whether cells the model ranks high on an axis carry more of the corresponding measured feature, cells were ranked within each sample by the signed SHAP^31^ contribution along that axis. The top 25% and the bottom 25% of cells within each sample were taken as the high- and low-groups. The measurement values underlying the high- and low-SHAP cumulative distributions were compared using a two-sided Mann–Whitney U test. External validation used an in-house esophageal cohort, comprising 18 tumor samples from 18 patients (one sample per patient, 144,292 cells), each with a whole-slide H&E image and DAPI imaging of the same section. The cohort was used only for evaluation and DAPI supplies only the observed measurement. A DAPI counterpart was pre-specified per feature: nuclear area, aspect ratio, and solidity are the same physical quantity measured on an independent segmentation, whereas chromatin density, coarseness, and pattern map to labeled scalar analogs (DAPI inverse mean, DAPI intensity standard deviation, and DAPI inverse skewness), and eosinophilic foci has no single-channel DAPI counterpart. DAPI measurements were z-scored using the paired cells of their own sample, matching the correction applied to the H&E targets, and the two are labeled in their different units.

#### Esophageal patient sample preparation

For independent validation we used 18 in-house tumor samples from 18 esophageal cancer patients, each contributing a paired whole-slide H&E image and DAPI imaging of the same physical section, prepared as follows. Tumor specimens were collected from patients with esophageal cancer who underwent chemoradiotherapy at The University of Texas MD Anderson Cancer Center (Houston, TX, USA). The study protocol was reviewed and approved by the MD Anderson Institutional Review Board, and all participants provided informed consent. Histological diagnoses were independently verified by experienced pathologists. Formalin-fixed, paraffin-embedded (FFPE) tissue blocks were sectioned at 5 µm on a rotary microtome, floated on a 40–42 °C water bath, mounted on Fisherbrand™ Superfrost™ Plus Microscope Slides compatible with the Akoya Phenocycler Fusion 2.0 instrument, and dried at 60 °C for 1 h prior to staining. Sections were deparaffinized in xylene, rehydrated through a graded ethanol series, and subjected to heat-induced epitope retrieval in pH 9 buffer (Akoya Biosciences) using BioGenex EZ-Retriever® at 98 °C for 20 min. Slides were blocked for 1 h at room temperature and stained overnight at 4 °C with the Akoya Biosciences IO60 antibody panel supplemented with seven additional in-house-conjugated, barcoded antibodies. The assay reporters were prepared, and the PhenoCycler-Fusion experimental run was performed using Akoya’s Fusion 1.0.6 software according to Akoya’s PhenoImager Fusion User Guide. Images were taken on the PhenoImager Fusion microscope using default settings. The best focus projections were computed, tiles were stitched, and cycles were registered using the DAPI channel in Fusion Processor software. The staining images (including DAPI) were exported as .QPTiff images for downstream analysis. After completion of all imaging cycles, coverslips were removed from the same physical sections and slides were washed in Histochoice to remove residual reporter oligonucleotides. Slides were stained with Mayer’s hematoxylin (1–5 min), blued in running tap water, counterstained with eosin Y (30 s–2 min), and coverslipped with a permanent mounting medium (Permount, Fisher Scientific). Whole-slide brightfield H&E images were acquired at 20× magnification on the PhenoCycler Fusion microscope using the brightfield mode, exported and used for further analysis.

### Morphology-based pseudotime inference

For diffusion map^51^ analysis, we utilized H&E images from skin and brain samples obtained from 10x Genomics (https://www.10xgenomics.com/datasets), together with two CRC patient samples^34^. Cell-level morphological embeddings were first extracted by Loki2 and used to construct a k-nearest neighbor graph, capturing local relationships between cells in the high-dimensional feature space. Diffusion maps were then computed by modeling transitions between neighboring cells as a Markov process on this graph, followed by eigen decomposition of the diffusion operator. To quantify the degree to which the learned diffusion manifold exhibits linear structure, we computed a linearity score based on the anisotropy of the two-dimensional diffusion map embedding. For each tissue, we selected the two diffusion components that showed the highest correlation with morphological pseudotime. We then calculated the covariance matrix of this point cloud and obtained its eigenvalues *λ*_1_ ≥ *λ*_2_. The linearity score was defined as:

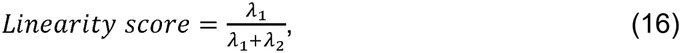

a value close to 1 indicates a highly linear structure, where variance is concentrated along a dominant axis, whereas lower values indicate a more isotropic distribution. For cross-tissue single-cell morphology–transcriptome similarity analysis, we utilized H&E images paired with single-cell-resolution spatial transcriptomics data across multiple organs, including skin, lung, colon, kidney, liver, ovary, breast, and brain, from 10x Genomics (https://www.10xgenomics.com/datasets), where the colon sample was the P1CRC sample previously used in single-cell morphology-to-transcriptome retrieval task. The Loki2 model was employed to extract cell-level morphological embeddings from histological images, while OmiCLIP was used to derive corresponding transcriptomic embeddings from gene expression profiles. Both embedding modalities were subsequently projected into a shared 512-dimensional latent space, following the procedure described in the single-cell morphology-to-transcriptome retrieval task. This dimensional alignment enables direct computation of cosine similarity between morphology-derived and transcriptome-derived representations, facilitating quantitative assessment of cross-modal concordance across tissues.

To infer cancer cell progression from morphological snapshots, we adapted Palantir^33^, a graph-based algorithm originally designed for scRNA-seq data. We posit that cell morphological embeddings from Loki2, like transcriptional profiles, lie on a low-dimensional manifold where phenotypic changes occur incrementally. High-dimensional embeddings from Loki2 define this morphological space. A nearest-neighbor graph was constructed in this space, with nodes representing individual tumor and epithelial cells and edges linking morphologically similar states. This graph approximates the continuous phenotypic manifold, capturing gradual transitions associated with cancer progression. Pseudotime inference was modeled as a stochastic process using a Markov chain on the neighbor graph. Edges were oriented by their distances from a user-defined root cell assuming unidirectional progression. The algorithm assigns pseudotime values to cells and identifies terminal states as outliers in the stationary distribution of the random walk, enabling quantitative ordering of cells along a morphological progression axis.

For the CRC analysis, one grade 1 conventional adenoma (TA/TVA) and two grade 2 microsatellite stable (MSS) tumor samples annotated as G1-Clone 1 and G2-Clone 2/Clone 3, based on the CNV clusters from the original study^34^ were examined. Loki2 was applied to the WSIs for nucleus segmentation and cell classification. Morphological embeddings for neoplastic cells were used for trajectory analysis. To capture global relationships among samples from various grades, partition-based graph abstraction (PAGA)^52^ was first applied. A nearest-neighbor graph was then constructed, initialized from the PAGA topology. A cell from G1-Clone 1 sample was designated as the root, and pseudotime was inferred across all samples. For the prostate cancer analysis, section ‘H2_5’ annotated with benign, transition state, and Gleason grade group 4 (GG4) cribriform regions from the original study^35^ was analyzed. A cell from the benign region was set as the root based on the spatial context and the UMAP embedding. The number of neighbors was set to 100 for constructing the nearest-neighbor graph, and the number of waypoint sampling = 2,000 was employed for stable pseudotime estimation. For pseudotime-associated gene analysis in the prostate cancer sample, we utilized gene expression profiles from paired Visium spatial transcriptomics data. Cell-level pseudotime were first inferred by Loki2 and subsequently aggregated to the spot level by averaging pseudotime across all cells mapped to each Visium spot. We then computed the Pearson correlation coefficient between gene expression and spot-level pseudotime for each gene, thereby quantifying the degree of association with the inferred pseudotime. Genes were stratified into three groups based on their correlation with pseudotime: Group 1 (absolute value < 0.2), Group 2 (absolute value 0.2–0.4), and Group 3 (absolute value > 0.4). Pathway enrichment analysis was subsequently performed within each category to elucidate functional programs linked to cellular progression along the inferred pseudotime.

### Single-cell morphology-to-transcriptome retrieval

To enable the prediction of single-cell transcriptomes directly from histology images, we developed a retrieval-based task rooted in contrastive learning. The model was trained and evaluated on the Visium HD colorectal carcinoma dataset (10x Genomics)^54^ comprising five distinct tissue samples obtained from three patients: ‘P1CRC’, ‘P2CRC’, ‘P3NAT’, ‘P5CRC’, ‘P5NAT’, where ‘Pn’ denotes the patient ID, ‘CRC’ denotes colorectal cancer, and ‘NAT’ denotes normal adjacent tissue. For all methods in this evaluation, only the paired H&E and Visium HD data from the ‘P1CRC’ sample were utilized for the cross-modal alignment training phase. Data from all other samples were reserved exclusively for inference and retrieval evaluation.

Histological images were segmented into nuclei using Loki2, yielding cell-level morphological embeddings denoted as *ϵ*_*I*_ ∈ ℝ^1280^. Corresponding transcriptomic profiles were aggregated to overlapping cells based on the Visium HD data (2 *μm* bins) and embedded using OmiCLIP11, producing transcriptomic embeddings *ϵ*_*T*_ ∈ ℝ^768^. A dual-encoder architecture was employed to align morphological and transcriptomic features through contrastive learning. To map the heterogeneous representations into a unified latent space, a single-layer linear projection was applied to each modality, defined as:

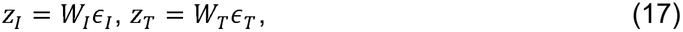

where *W*_*I*_ ∈ ℝ512×1280 and *W*_*T*_ ∈ ℝ512×768 are learnable projection matrices, and *z*_*I*_, *z*_*T*_ ∈ ℝ512 are the resulting projected morphology and transcriptomic embeddings. Bias parameters were omitted to enforce angular alignment between the projected vectors. During training, both the morphology and transcriptomic encoders were frozen, and only the projection layers were optimized. We employed a symmetric cross-entropy contrastive loss designed to maximize the cosine similarity sim(·,·) between matched morphology–expression pairs defined as:

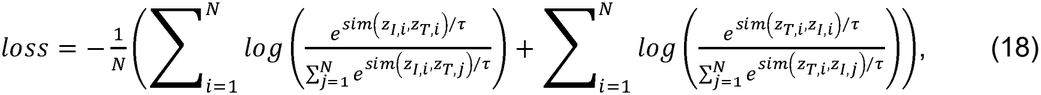

where *N* is the batch size, *τ* is the temperature parameter controlling distribution sharpness, *z*_*I,i*_ and *z*_*T,i*_ represents the projected morphology and transcriptomic embeddings for the *i*-th cell. Training was conducted using the AdamW optimizer with a learning rate of 5 × 10^™4^ for 20 epochs with a batch size of *N* = 4,096.

Transcriptomic prediction was achieved by projecting Loki2 morphology embeddings encoded from query histology images into the shared latent space and retrieving the nearest neighbor RNA expression profiles from a reference database. Two inference modes were implemented to support distinct use cases. In out-of-sample retrieval, predictions are made by comparing query cells to a reference atlas or an external scRNA-seq database from the same tissue type. Let *z*_*I*,query_ ∈ ℝ^512^ denote the projected morphology embeddings of a query cell. For each cell type *C*, let *D*_*C*_ = {*z* ∈ ℝ^512^} denote the set of projected transcriptomic embeddings of all reference cells annotated as type *C*. A two-step hierarchical retrieval strategy was applied to ensure biologically coherent matches. First, the most probable cell type *C*_best_ was selected by maximizing the mean cosine similarity between the query embeddings and all transcriptomic embeddings belonging to that type:

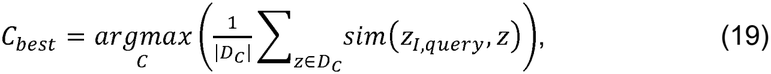

where |*D*_*C*_| is the number of reference cells in class *C*. Next, the predicted transcriptomic embeddings 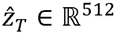 were retrieved by selecting the single cell within the chosen type-specific subset 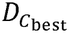 that maximizes similarity to the query:

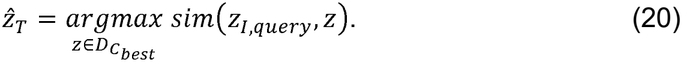

In in-sample retrieval, used when a large H&E slide is only partially profiled by Visium HD, regions of the H&E image lacking RNA-seq data can directly query the profiled RNA-seq data from the same tissue section. Let *D*_in-sample_ denote the set of transcriptomic embeddings from the sequenced regions of the same specimen. The predicted scRNA-seq profile is retrieved via:

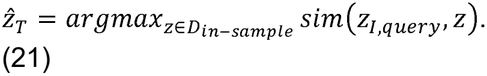

For GHIST and iStar models, training was performed using paired H&E histology and Visium HD transcriptomic data from the sample P1CRC, with cell-type annotations withheld to ensure fair comparison with Loki2. GHIST was trained using the default architecture and hyperparameters specified in its official repository^23^ with nuclei segmentation masks, identical to those used in the Loki2 pipeline, provided as input. Because both GHIST and iStar are optimized for predicting highly variable genes, we restricted training to genes exhibiting strong spatial patterns, defined as those expressed in at least 1,000 cells and having Moran’s I > 0.01, resulting in a final panel of 915 genes used for training and inference. iStar was deployed using its recommended default configuration from the official documentation^36^, trained on H&E images paired with Visium HD data (8 bins) from P1CRC, with an output pixel size of 0.5 to enable high-resolution gene-expression mapping. To facilitate cell-level performance evaluation and direct comparison with other methods, the predicted superpixel-level gene expression values were aggregated to single-cell resolution using the same segmentation masks. After training on P1CRC, both GHIST and iStar were used to generate spatial gene-expression predictions for the independent test samples (P2CRC, P3NAT, P5CRC, and P5NAT).

For the sample ‘P1CRC’, both in-sample and out-of-sample retrieval modes were applied. For all other samples, only the out-of-sample mode was utilized to assess generalization capability. For statistical evaluation, the Visium HD data were used as the ground truth. The ground truth cell types were determined by CellTypist^55^, utilizing reference scRNA-seq annotations from the original study^54^. Prediction accuracy was quantified using Pearson’s correlation coefficient and Spearman’s rank correlation coefficient to assess gene expression fidelity in a cell. Structural similarity index measure (SSIM), which ranges from -1 to 1, was computed to evaluate the preservation of spatial expression patterns of a gene, where we considered the gene distribution in spatial as an image. For two images and :

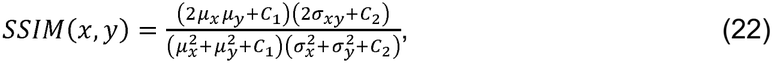

where: *u*_*x*_ and *u*_*y*_ are the mean intensities of images *x* and *y*, 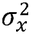 and 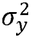 are the variances of *x* and *y, σ*_*xy*_ is the covariance between *x* and *y, C*_1_ and *C*_2_ are small constants to stabilize the division when the denominators are close to zero.

For the platform-dependency analysis, we used the same workflow in which the model was trained on paired H&E images and Xenium spatial transcriptomics data instead of Visium HD. All other settings were kept unchanged, including the training and test samples, reference scRNA-seq dataset, and evaluation metrics. This design enables systematic assessment of the extent to which model performance depends on the underlying spatial transcriptomics platform. For the reference scRNA-seq dependency analysis, we used an external reference scRNA-seq dataset^45^ for retrieval. The pretrained model using Visium HD data was applied to the same test samples without additional retraining, while only varying the used scRNA-seq data for retrieval. This setup isolates the influence of reference atlas selection on downstream performance. We further constructed reference variants with different mixing ratios between the original and external atlases to evaluate the stability of downstream results under varying reference conditions. These included the original atlas only, 80% original plus 20% external, 60% original plus 40% external, 20% original plus 80% external, and the external atlas only.

For the biomarker discovery study, the pretrained model using Visium HD data was applied to H&E images from 100 CRC patients in the PLCO cohort without additional retraining, including 50 low-grade and 50 high-grade patients. Retrieved transcriptomic profiles were derived from an external single-cell RNA-seq reference dataset^38^. We first identified the top three differentially expressed genes in the retrieved tumor cells by comparing low- and high-grade patients. We next tested this three-gene signature in an independent TCGA colorectal cancer cohort (n = 151), derived from the rectal adenocarcinoma dataset. For each patient, a gene-set score was calculated as the mean expression of the selected genes. Patients were then stratified into high- and low-score groups using the median gene-set score as the cutoff. Statistical significance between survival curves was assessed using a two-sided log-rank test.

### Morphology-molecular coupling analysis

Analyses used fifteen 10x Genomics Visium HD sections with paired H&E images (three breast cancer, three colon cancer, two normal colon, three lung cancer, two lymph node, one kidney, one pancreas). Nuclei were detected on each H&E image with Loki2, which returns a 1,280-dimensional morphology embedding per nucleus. Genes were selected within each section: non-zero in at least 3% of cells and in at least 20 cells, then normalized to 10,000 counts per cell (CP10k) and log1p-transformed, then required to have mean log1p(CP10k) ≥ 0.3.

For the gene-property analysis, the mean-centered embedding of each section was decomposed by PCA, retaining the components carrying 90% of the variance. Each morphology component was cut into 20 equal-frequency bins *B*_*k*_ and each gene into four states *S*_*g*_, namely ‘not detected’ plus tertiles of its positive values, and mutual information was computed in bits from James–Stein shrinkage entropies^39^,

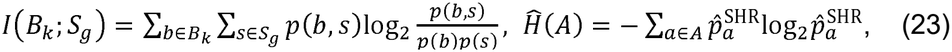

where 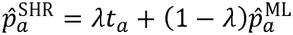 shrinks the maximum-likelihood cell frequencies 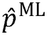 of a discrete distribution toward the uniform target *t*_*a*_ = 1/|*A*|, with the shrinkage intensity *λ* estimated analytically. Each gene was scored by its maximum over the components of its section. Scores were pooled by taking the median across the sections. The genes are split into five equal-sized groups (G1 highest, G5 lowest) by score.

The linear-probe analysis used the Loki2 1,280-dimensional embedding as predictors, with training-fold column means subtracted, and log1p(CP10k) as the response. Cross-validation used five contiguous spatial bands: centered coordinates were decomposed by singular value decomposition (SVD), cells were projected onto the first right singular vector, and the rank- ordered projections were split into five bands of 2,000 cells. Ridge regression was solved in closed form from one SVD per training block, serving all genes and all penalties,

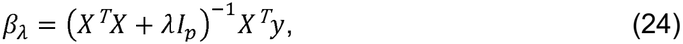

Penalties were selected inside the training data: each 8,000-cell training set was split into four inner spatial bands by the same algorithm, inner held-out Pearson correlations were summed over the inner folds for each gene and each of eleven candidate penalties *λ* ∈ {10^0^, 10^1^, …, 10^10^ }, and the maximizing value was taken; the penalty finally used was the median grid index of the five outer folds’ choices. A gene’s probe score is the held-out Spearman correlation between its predicted and observed expression, averaged with equal weight over the five outer folds.

Gene-set function naming followed the protocol of Hu et al.^53^ that each group of genes (G1 to G5) was submitted to a large language model (Claude Opus 5). Two properties were computed: cell type numbers and cell type specificity, the Tau specificity index over a partition of the section’s cells into groups, computed from unweighted per-group CP10k means *m*_*gi*_ over groups.

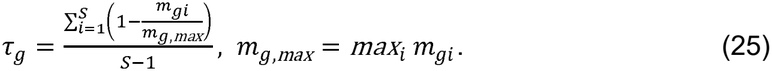

### *In silico* immunostaining from standard histology

H&E WSIs were processed using Loki2 to obtain single-cell representations. For each WSI, Loki2 produced (i) cell-level embeddings and (ii) instance segmentation masks for cell nuclei. All cellular morphology embeddings were normalized by *ℓ*_2_-norm prior to downstream modeling. For an embedding vector *z*_*k*_ ∈ ℝ^*d*^ corresponding to a cell *k*, we computed the normalized embedding 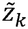 by 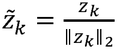. For the HEMIT dataset43, the corresponding IHC images were pre-aligned to the H&E WSIs by the data provider. For the CyCIF dataset (core_002)^41^, the mIF images were manually registered to the corresponding H&E WSIs. Let *T* denote the spatial transformation mapping coordinates from the H&E image domain Ω_*H&E*_ to the IF domain Ω_*IF*_; then, for each pixel *p* ∈ Ω_*H&E*_, the aligned IF signal was registered at *T*(*p*). Following registration, paired H&E and IF images were cropped into patches of size 2048 × 2048 pixels for all subsequent analyses.

To capture proteins that localize to the cell membrane, we expanded nuclear masks into approximate whole-cell masks using a marker-based watershed algorithm with a geodesic distance constraint. Let *G* ∈ ℝ^*H*×*W*×*C*^ denote the multi-channel IF patch used as guidance, and let:

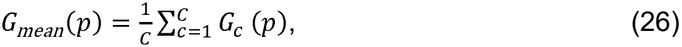

be the mean intensity across channels at pixel *p*. We define an inverted intensity surface

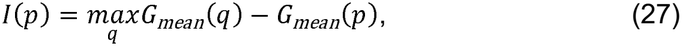

where the maximization over *q. q* is taken across all pixels in the patch and *I*(*p*) serves as the watershed’s topographic surface. Let *M*_*nuc*_ be the binary nuclear instance mask (markers) produced by Loki2 and let *d*_geo_(*p, M*_*nuc*_) denote the geodesic distance from pixel *p* to the nearest nuclear pixel along the intensity surface *I*. A marker-based watershed is then applied to *I*, seeded by *Mnuc*, and constrained to pixels satisfying *d*_*geo*_(*p, M*_*nuc*_) ≤ *d*_*max*_, where *d*_*max*_ is the maximum allowed geodesic expansion (default 20 pixels). Last, the watershed segmentation yields a label map *L*: {1, …, *H*} × {1, …, *W*} → {0, 1, …, *K*}, where *L*(*p*) = *k* indicates that pixel *p* is assigned to cell *k*, and *L*(*p*) = 0 denotes background. For each aligned IF tile, let *X* ∈ ℝ*H*×*W*×*C* denote the IF intensity image, where *X*_*c*_(*p*) is the intensity at pixel *p* in channel *c*. Given the cell label map *L*, the set of pixels belonging to cell *k* is *S*_*k*_ = {*p* | *L*(*p*) = *k*}. For a target protein channel *c*, we first compute the mean raw intensity for cell *k* as:

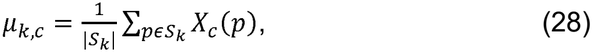

where |*S*_*k*_| denotes the number of pixels assigned to cell *k*. To stabilize variance and reduce the influence of extreme values, we use a log-transformed per-cell intensity as the target protein expression:

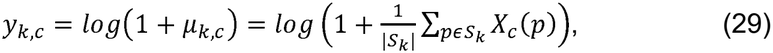

These scalar values *y*_*k,c*_ serve as the regression targets for predicting protein expression from H&E-derived cell embeddings.

To benchmark the prediction models, for each cell *k*, let 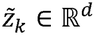 denote the *ℓ*_2_-normalized embedding obtained from the H&E image. For each protein channel *c*, we consider a scalar regression function:

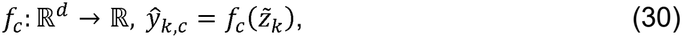

where *ŷ*_*k,c*_ is the predicted log-transformed protein expression for cell *k*. We benchmarked our method against several standard regressors trained independently for each protein channel *c* including: LR, SVR with radial basis function (RBF) kernel, MLP regressor with two hidden layers (256 and 128 units, ReLU activations, early stopping), LightGBM regressor (learning rate 0.05, num_leaves = 64, feature_fraction = 0.8, bagging_fraction = 0.8, early stopping on a validation set), and ROSIE (using released model, default parameters). Each model was trained using per-cell pairs 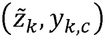. To assess generalization performance and select hyperparameters, we performed five-fold cross-validation, partitioning patches into five non-overlapping folds and using four folds for training and one for validation in each iteration.

For independent validation on the TCGA and PLCO cohorts, we directly applied a pretrained LightGBM model that previously trained on CRC patient sample to predict the PanCK protein marker, to the Loki2-derived cell embeddings without any additional finetuning. This evaluation was conducted on a total of 10 CRC samples, comprising 5 samples from the TCGA cohort and 5 samples from the PLCO cohort. This approach enables a rigorous assessment of model generalizability across external datasets. To facilitate quantitative comparison across samples with varying signal distributions, we performed within-sample standardization by computing Z-scores of the predicted PanCK values. Specifically, for each sample, predictions were normalized by subtracting the sample-specific median and dividing by the corresponding standard deviation.

#### Evaluation metrics

Mean absolute error (MAE): Let *y* = (*y*_1_, …, *y*_*n*_) denote the ground-truth per-cell protein expressions and ŷ= (ŷ_1_, …, ŷ_*n*_) the corresponding predictions for a given protein channel. We report multiple complementary metrics.

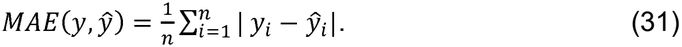

SSIM was computed using the same formulation as in the single-cell morphology-to-transcriptome retrieval analysis. All metrics were computed on held-out test cells for each protein channel independently.

### Cell-level multiple instance learning

This task leveraged the TCGA cohort, including KIRC, COAD, and BLCA, to construct a weakly supervised learning task. For each cancer type, we select 120 patients for tumor stage, using their histology image from each of the patients. To mitigate class imbalance, we performed down-sampling to obtain 60 paired early- and late-stage samples. This yielded a balanced 1:1 patient ratio for subsequent stage classification. We define the tumor stage clinical task as carcinoma stage prediction, a binary classification task stratifying patients into early (I/II) versus late (III/IV) stages, denoted as:

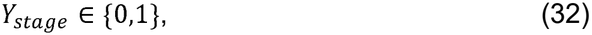

where 1 represents late stage, and 0 denotes early stage.

We leverage an MIL to analyze the Loki2-generated dataset 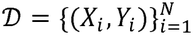 for these objectives. Given dataset 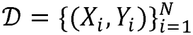 for each of *N* patients, where 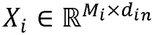 represents the M_*i*_ single-cell morphology embeddings with dimension *d*_*in*_ = 1,280 derived from the WSI of patient *i*, and *Yi* denotes the corresponding clinical label. Data loading and alignment are performed by matching the unique patient identifier between the down-sampled embeddings (averaging 10,000 single-cell embeddings per patient) and clinical metadata.

To map the unordered set of single-cell embeddings to a patient-level clinical prediction, we implemented a gated MIL consisting of three distinct components: a non-linear feature projection head, a gated attention module, and a global aggregation classifier. For each patient *i*, the WSI is processed to yield a bag of instances 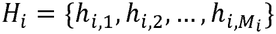. Each instance embedding 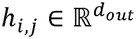 is extracted from the raw input *x*_*i,j*_ via a four-layer perceptron, formally defined as a composition of non-linear transformations:

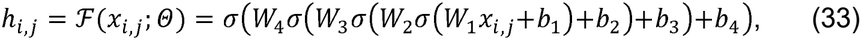

where *d*_*out*_ = 1,280, *σ*(·) denotes the ReLU activation function, and 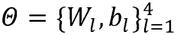 represents the learnable parameters, where *W* is weight and *b* is bias. The layer dimensions follow a bottleneck architecture defined by the sequence *d* = [1280, 1024, 768, 512, 1280], such that 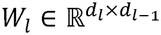 and 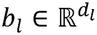.

To prevent overfitting and ensure training stability, we apply distinct regularization techniques: high-rate dropout percentage *p* (*p* = 0.6) is utilized in the initial layers, while batch normalization is applied to the penultimate layer (512-dim) prior to the final projection. To capture the heterogeneity of the tumor microenvironment, we employ a learnable gated attention mechanism that assigns an importance score *a*_*i,j*_ to the *j*-th instance of patient *i*, allowing the model to focus on diagnostic regions (e.g., invasive tumor area) while suppressing background noise. The attention score is computed as:

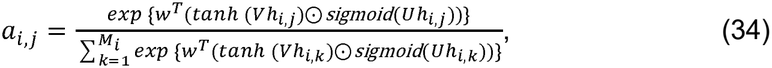

where 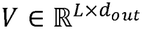 and 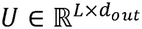 are weight matrices, and *w* ∈ ℝ^*L*×1^ is a weight vector, with attention dimension *L* = 128. The element-wise product (⊙) of the hyperbolic tangent and sigmoid activations act as a gating mechanism, enabling non-linear interactions between features to modulate the attention distribution. The slide-level latent representation *Z*_*i*_ is obtained via the attention-weighted aggregation of all cellular instances:

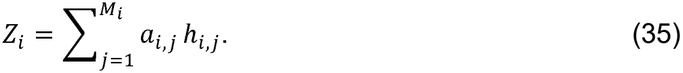

Finally, a linear classifier *g*_*ϕ*_ maps the aggregated feature Z_i_ to a predictive logit score. We utilized the binary cross entropy (BCE) with a logits loss function for optimization, which combines a Sigmoid layer and the BCE loss into a single class for improved numerical stability. For inference, the final probability Ŷ_i_ is derived as:

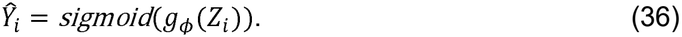

Probabilities are clamped to the interval [10^™7^, 1 ™ 10^™7^] to prevent numerical underflow/overflow during evaluation.

To rigorously evaluate model generalization and mitigate class imbalance, we employ a stratified *K*-fold cross-validation strategy with *K* = 5. The dataset *D* is partitioned into *K* disjoint subsets {*D*_1_, …, *D*_*K*_}. To prevent data leakage, the partition is performed at the patient level. We utilize a balanced fold assignment algorithm to ensure that the ratio of positive to negative samples in each fold approximates the global population ratio:

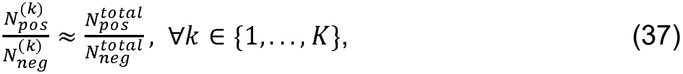

where 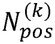 and 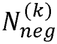 denote the number of positive and negative cases in the *k*-th fold, respectively. Global population ratio of each task selected as 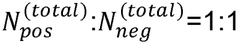, where the total amount of patients for the stage task is 120. For each iteration *k, D*_*k*_ serves as the test while the remaining folds are further subdivided into training (80%) and validation sets (20%) for the model’s early stop mechanism. To compare fairly, we obtain the average results for comparison after training five times with different seeds.

For slide-level evaluation, we compared Loki2 with representative foundation models, including PLIP, UNI, and GigaPath. For UNI, we followed the default pipeline to crop WSIs into patches and directly extract slide-level features. For PLIP, we applied the default preprocessing pipeline, which resizes WSIs to generate slide-level embeddings. For GigaPath, we followed the default pipeline to crop WSIs into patches and subsequently applied mean pooling across patch embeddings to obtain a single slide-level features. These slide-level features were used as inputs for RF classifiers configured with 500 trees and balanced class weights to perform binary classification of each cancer stages. To ensure a fair and rigorous comparison, the dataset partitioning and training protocols were kept strictly identical to our approach, utilizing the same stratified five-fold cross-validation strategy for the binary prediction task.

#### Evaluation metrics

Receiver operating characteristic (ROC): To assess the trade-off between sensitivity and specificity at various threshold settings, we compute the area under the ROC curve (AUC). The ROC curve plots the true positive rate (TPR) against the false positive rate (FPR), defined as:

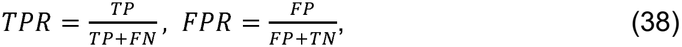

where TP, FN, FP, and denote true positives, false negatives, false positives, and true negatives, respectively.

Precision-recall (PR): We also evaluated the average precision (AP), where Precision and recall are defined as:

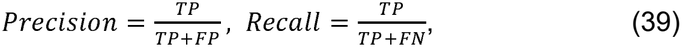

The summarizes the curve as the weighted mean of precisions achieved at each threshold, providing a robust metric for minority class detection.

## Data Availability

The histology images of the TCGA dataset used for the cell type annotation task, morphology-to-transcriptome retrieval task, *in silico* immunostaining task, and cell-level whole-slide clinical prediction are available from the NIH Genomic Data Commons (http://portal.gdc.cancer.gov). The human dorsolateral prefrontal cortex H&E images used for the model validation can be found at https://www.nature.com/articles/s41593-020-00787-0^20^. The human colorectal cancer dataset, including H&E image, Visium-HD, Xenium, and scRNA-seq data used for the morphology-to-transcriptome retrieval task can be found at: https://doi.org/10.1101/2024.06.04.597233^54^. The external colorectal cancer scRNA-seq data used for morphology-to-transcriptome retrieval task can be found at: https://www.cell.com/cancer-cell/fulltext/S1535-6108(24)00234-4^38^ and https://www.sciencedirect.com/science/article/pii/S0092867421009454^45^. The histology images of the PLCO dataset used for the morphology-to-transcriptome retrieval task and *in silico* immunostaining task are available from the NIH Cancer Data Access System (https://cdas.cancer.gov/datasets/plco/). The H&E images and paired single-cell-resolution spatial transcriptomics data of skin, lung, kidney, liver, ovary, breast, colon, and brain samples used for cross-tissue single-cell morphology–transcriptome similarity analysis are available from 10x Genomics: (https://www.10xgenomics.com/datasets). The colorectal cancer data used for the morphological pseudotime inference task can be found at: https://doi.org/10.1016/j.cell.2023.11.006^34^. The prostate cancer data used for the morphological pseudotime inference task can be found at: https://doi.org/10.1038/s41586-022-05023-2^35^. The paired CRC CyCIF and H&E (core 002) used for the *in silico* immunostaining task were downloaded from Amazon Web Services S3 (s3://lin-2021-crc-atlas/data/) from GitHub https://github.com/labsyspharm/CRC_atlas_2022^41^. The paired colon cancer IHC and H&E data used for the *in silico* immunostaining task were downloaded from https://github.com/BianChang/HEMIT-DATASET^43^.

## Code Availability

Loki2 is implemented in Python and is available via https://github.com/GuangyuWangLab2021/pathweb/. The source code is in the Supplementary

File, the pretrained model weights is available via https://drive.google.com/drive/folders/1l1f8A3qcC-ChjCl5nST4IC5Dr2ZbNpfb?usp=sharing, and the demonstration data for the tutorial is available via https://drive.google.com/drive/folders/19qzd8jTZBQ57a3N5xbiZFSGZJNpIz8NQ?usp=sharing (source code and pretrained model weights will be released on GitHub and Hugging Face after the manuscript is accepted).

